# Overwintering performance of juvenile temperate estuarine fish

**DOI:** 10.1101/2024.03.28.587281

**Authors:** Clara Bellotto, Ashley M. Fowler, David J. Booth

## Abstract

Estuaries, rich in biodiversity and economically valuable species, are increasingly threatened by climate change-induced factors that challenge fish resilience and survival. This study compared the performance of estuarine fishes between water temperatures reflecting two scenarios: current Sydney winters (16°C) and future winters under climate change (20°C), and at two food levels, for three estuarine fish species (eastern fortescue, *Centropogon australis*, common silverbiddy, *Gerres subfasciatus,* and eastern striped trumpeter, *Pelates sexlineatus*) Overall, as expected from metabolic theory, fish performance was generally higher at higher temperatures, with growth rates higher at 20°C for *G. subfasciatus* and *C. australis.* Bite rates and aerobic scope were generally higher at the higher temperature for all species. *G. subfasciatus* and *P. sexlineatus* exhibited increased escape responses at 20°C, with *P. sexlineatus* also showing greater boldness. Boldness was positively associated with bite rates in *P. sexlineatus*, potentially indicating foraging advantages under future warming for this species. The order of temperature treatment (20°C then 16°C, vs 16°C then 20°C) affected boldness for *G. subfasciatus* and growth rate, total length, bite rate and burst speed for *P. sexlineatus*.

Contrary to expectations, food had no effect on fish performance either directly or interacting with temperature, and all three species generally performed better at 20°C than 16°C, suggesting this study was conducted below the species’ thermal optima. Future climate change may therefore favour temperate estuarine fishes at winter temperatures, with potential benefits differing among these species.

## 1. Introduction

Australia’s oceans are a hotspot of marine biodiversity and warming (Wernberg *et al*., 2011). Globally, south-eastern Australia has experienced one of the fastest increases in mean ocean temperature, which has increased at approximately four times the global average (0.023°C year^-1^, (Ridgway, 2007). This increase is mainly driven by strengthening of the poleward flowing East Australian Current (Ridgway, 2007; Figueira and Booth, 2010). In addition to oceans, it was recently estimated that Australian estuaries are warming at an alarming rate of 0.2 °C year^−1^ (Scanes *et al*., 2020). Despite these significant changes, the response of estuarine systems to climate change is is under-researched with respect to coastal habitats (O’Brien *et al*., 2016; Khojasteh *et al*., 2021).

A critical environmental variable is ambient temperature; as fish are poikilotherms, the water temperature affects fish physiology and behaviour where their body temperature conforms to the surrounding water temperatures (Angilletta Jr *et al*., 2002; Booth *et al*., 2011). Temperature possesses a direct thermodynamic effect on biochemical reaction rates (Little *et al*., 2020). Consequentially, temperature influences all aspects of physiology and behaviour in ectotherms; these include sensory inputs, foraging abilities, locomotion, growth and immune function (Matear *et al*., 2000; Du Pontavice *et al*., 2020). The key drivers of this process are fish’s relatively low metabolic rate, the lack of insulation, and the high specific heat of the water (Beitinger *et al*., 2000).

As ectotherms are exposed to a broad temperature range, the relationship between body temperature and physiological performance is expected to be a bell-shaped function, where performance is maximum at intermediate temperatures below which performance increases with temperature and after which performance decreases with further increases in temperature. To understand the relationship between the thermal environment and organismal performance, biologists rely on several characteristics of performance functions. The thermal optimum is the body temperature at which performance peaks. Critical thermal limits define the temperature range necessary to sustain basic performance (Angilletta Jr *et al*., 2002). If environmental temperature shifts outside this range fish physiological performance drops rapidly and mortality risk increases greatly (Westhoff and Paukert, 2014). However, performance can vary across species and individuals in a given environment based on plasticity during development (developmental plasticity), during the juvenile or adult phase (reversible acclimation) or by previous generations (transgenerational plasticity) (Gilchrist, 1995; Burton and Metcalfe, 2014). Because of the natural fluctuation of environmental conditions in estuaries (Scanes *et al*., 2020), we might expect species performance to be similar across a wide temperature range, indicating considerable plasticity (Gunderson and Stillman, 2015).

Due to the significant impact of body temperature on performance, it is unsurprising that ectotherms have evolved various mechanisms to respond to the heterogeneity of the thermal environment (Little *et al*., 2020). Individuals have been shown to use both behaviour (e.g., avoidance/preference) and physiology (e.g., acclimatisation) to regulate their body temperature within a narrower range than the range of ambient temperatures. This process that assists in dealing with spatial and thermal heterogeneity in the thermal environment is called thermoregulation (Angilletta Jr *et al*., 2002). However, optimising the performance function by change in behaviours may be more effective than relying solely on thermoregulation, which can increase mortality risk, require more significant energy expenditure, or lead to missed opportunities (Angilletta Jr *et al*., 2002).

Within an individual’s lifespan, performance functions can be modified by acclimatising to environmental temperatures via reversible acclimation (Little *et al*., 2020). This can occur through various cellular processes such as allozyme expression or cell membrane changes. Over a few generations, the thermal environment can affect the evolution of an organism’s performance functions. Thus, natural selection can modify essential parameters of an organism’s performance functions and the ability of this organism to acclimate and adapt to changes in these parameters in response to the thermal environment (Kingsolver and Huey, 1998). The temperature that fish experience during their larval stage affects its plasticity and can profoundly influence fish fitness and, thus, future stocks abundances and composition (Burton and Metcalfe, 2014). Therefore, looking at how higher temperatures will impact fish at their younger life stages is imperative, given the importance of early life history stages in determining adult abundance (recruitment limitation hypothesis: Doherty & Fowler 1994). Therefore, this study was conducted with juvenile fish and winter conditions (recruitment season). Plus, at this early life stage, selection likely had little time to decrease the variation within a population, increasing the possibility of detecting performance effects on fitness (Metcalfe *et al*., 2016).

A fish’s capacity to cope with environmental change includes physiological and behavioural responses (Alfonso *et al*., 2021). This study primarily focuses on temperature, because of its profound effects on ectotherms’ physiological and behavioural performance (Angilletta Jr *et al*., 2002). Other factors such as food availability also affect fish performance (Sinha *et al*., 2012), and this has rarely been studied especially as it interacts with temperature (Metcalfe *et al*., 2016). As such, this study investigated the effects of temperature and food regimes (usually limited during winter (Kuzuhara *et al*., 2019)) on a suite of performance metrics (metabolic rate, growth, burst speed, foraging behaviour, boldness, shelter and escape response).

This study aimed to better understand the physiological and behavioural differences that may be expected under warmer winter temperatures associated with climate change for juvenile temperate estuarine fish. This may aid in predicting the consequences for future estuarine ecosystems, fisheries, and aquaculture (Alfonso *et al*., 2021). Specifically, we aimed to evaluate performance of common juvenile estuarine fishes (the eastern fortescue, *Centropogon australis*, the common silverbiddy, *Gerres subfasciatus*, and the eastern striped trumpeter *Pelates sexlineatus*) across various metrics at different temperatures and food regimes that reflect the Sydney region’s current and forecasted winter conditions. The experimental temperatures chosen were (1) current mean Sydney winter water temperature (16°C,) and (2) the climate-change forecasted water temperature in the Sydney region (20°C) (Booth and Sear, 2018). Additional temperatures were not investigated because they are not required as we did not seek to develop performance curves but rather compare two key temperature scenarios as outlined above. Furthermore, climate change induces changes in food resources which might be lower for some functional groups, with food scarcity common in winter (Diamond *et al*., 2017; Kim *et al*., 2019). We expected that fish population performance in general would be lower at lower temperatures and lower food levels and the extent of this effect would vary across species. We also predicted that some performance metrics (growth, total length, burst speed, bite rate, aerobic scope, and boldness) would be positively related to each other while others will be negatively related (escape and shelter response with boldness).

## 2. Materials and Methods

### 2.1 Ethical statement

All practices in this study were conducted following the NSW Department of Primary Industries scientific collection permit (Permit No: F94/696(A)-9.0) and the University of Technology Sydney Animal Ethics protocols (Permit No: ETH21-6609).

### 2.2 Fish species and collection

Juvenile fish were collected in Careel Bay, New South Wales, located in south-eastern Australia (33° 3’ 02.8’’ S; 151° 1’ 25.2’’ E) by towing a seine net (15 m x 2 m; mesh size 16 mm) parallel to the shoreline for approximately 20 m over sand, *Zostera* and *Posidonia* seagrass beds (water depth < 1.70 m). The catch was emptied into a 40 L water tub, where it was sorted, and non-targeted individuals were immediately released. Once an adequate sample size was reached for the species of interest, these fish were transported to aquarium facilities at the University of Technology Sydney (UTS).

This study included three estuarine temperate species: the eastern fortescue, *Centropogon australis*, *Scorpaenidae* (n = 20), the common silverbiddy, *Gerres* subfasciatus, *Gerreidae* (n = 37), and the eastern striped trumpeter, *Pelates sexlineatus*, *Tetrapontidae* (n = 39). These species were selected as, at collection time, individuals were available in sufficient sample size.

### 2.2 Laboratory husbandry and acclimation

Captured individuals were initially housed for two days in groups (n = 10-15 fish) in 40 L tanks at ambient Sydney estuary temperature at the time of capture (16°C-20°C). Fish were then separated and housed in individual 15 L tanks filled with 35 ppt natural seawater from holding tanks in the UTS facilities. Fish were acclimatised to the desired temperatures (16°C or 20°C) by a 0.5°C daily temperature change using 25 W heaters to ensure that acclimatisation occurred at an appropriate rate while minimising fish stress (O’Brien *et al*., 2022).

The acclimatisation period ranged from 1 to 7 days, depending on the initial temperature detected at the collection site and the treatment target temperature targeted temperature for the treatment as fish species were collected in different months (August/16°C for *C. australis*, October/18°C for *G. subfasciatus* and December/20°C for *P.sexlineatus*). The laboratory room where the experiment was conducted was at 14°C air temperature, lighting on a 12 h light:12 h dark cycle. The fish were fed New Life Spectrum® fish pellets (high-quality protein source, 0.5 mm diameter) two times a day according to the treatment food regime (see below). Two polyvinyl chloride (PVC) pipes were placed into the tanks to provide shelter, and cut-out pieces of black foam board were placed to partially cover the tank’s exterior to minimise external disturbance (O’Connor and Booth, 2021). Removal of debris (uneaten food and faeces) by siphoning, and a 30% water change, was conducted daily to maintain water quality. The water removed through this process was replaced by water pre-acclimated to the experimental temperature. Water pH, dissolved oxygen, and salinity were measured with a multi-sensor probe every two days, while water temperature was assessed daily to ensure treatment temperatures were maintained. Fish health and behaviour were visually appraised and recorded daily, as per Djurichkovic *et al*. (2019).

The fish were housed in the laboratory for approximately nine weeks, and at the end of the experimental period, all fish were euthanised with ice slurry (Lidster *et al*., 2017).

### 2.3 Experimental protocols

Fish performance was evaluated using a broad range of metrics (metabolic rate, growth, burst speed, foraging behaviour, boldness, shelter and escape response). Fish growth rate and metabolic rate are important physiological metrics as they reflect the amount of internal energy possessed by the organism (Clark *et al*., 2013; O’Connor and Booth, 2021). Bite rate, burst speed, shelter response, boldness and escape response are part of important behaviours that reflect fish performance within the environmental and in the social group (White *et al*., 2013). In addition, otolith (ear stones) increments can provide valuable information on fish historical record of environmental conditions as ontogenetic or environmental changes are reflected into changes in increment depositions (Payan *et al*., 2004). Otolith increment width is frequently used as a proxy of somatic growth whereby fish with higher growth rates present wider increments (Molony and Choat, 1990; Labropoulou and Papaconstantinou, 2000; Megalofonou, 2006). The exact formulation of each metric is detailed in Section 2.4.

Experimental protocols are described in Supplementary Figure 1. Fish length and mass were measured, and fish were randomly assigned to a treatment divided into groups and housed in individual tanks. If different size classes were present, these were balanced across groups using a random stratified approach.

For the *G. subfasciatus* and the *P. sexlineatus* experiments, half of the fish experienced low food regimes, and the other half experienced high food regimes. Fish in high food treatment were fed 1% of fish weight twice a day in fish pellets and fish in low food treatment were fed 0.5% of fish weight twice a day in fish pellets as per Donelson *et al*. (2010); the normal feeding ratio for fish is approximately 0.7% of their body weight in dry food (Héroux and Magnan, 1996). The food regime was maintained throughout the whole experiment. However, in the *C. australis* experiment, food regimes were not evaluated due to an insufficient number of individuals. Fish were instead divided into two groups based solely on temperatures and were fed *ad libitum*.

All groups experienced two temperatures (16°C and 20°C) but in a different order to account for the potential confounding effects of pre-acclimation to the existing wild temp. Secondly, carryover effects from one temperature treatment might influence responses to the subsequent temperature treatment. For instance, if the cold treatment permanently alters physiology or performance, it could hinder or prevent the full expression of responses to the warm temperature treatment (McArley *et al*., 2017; Kim *et al*., 2019). One group was exposed to temperatures in descending order (from 20°C to 16°C), while the other group was exposed to temperatures in ascending order (from 16°C to 20°C).

Once the required initial temperature was reached (16°C or 20°C), total length and wet mass were measured. The fish were then exposed for ten days to the corresponding temperature treatment. After these ten days, the performance metrics were measured once. Afterwards, the fish were acclimated to the final temperature (20°C or 16°C). Once the individuals reached the required temperature, the fish’s total length and wet mass were measured again. The fish were then exposed for ten days to the corresponding temperature treatment. After these ten days, the performance metrics were again measured. The fish were then euthanised and dissected. Fish otoliths (ear stones) were extracted and analysed.

It took approximately three days to assess fish behaviour and three days to measure metabolic rate. The order in which individual fish were selected to be tested was randomised. For behavioural tests, fish were transferred into a pre-acclimated test tank with a gridded background (0.5 cm grids), and observations were filmed using a GoPro® Hero 6 located in front and at the same level as the base of the test tank. Videos were recorded at 120 frames per second. Fish were acclimated for 10 minutes in the test tank before proceeding with the different behavioural tests, which had an observation period of 3 minutes, with a further acclimation time of 2 minutes between tests as per Djurichkovic *et al*. (2019).

### 2.4 Fish performance metrics

**(a) Fish growth** was measured as the change in fish total length and wet mass (O’Connor and Booth, 2021). Instantaneous growth rate (*G_INST_*) was used to estimate somatic growth rate. This was calculated using the mass of the individual at the beginning (*M_1_*) and at the end (*M_2_*) of the experimental period (*t*, number of days), following equation 1 (Booth *et al*., 2014):

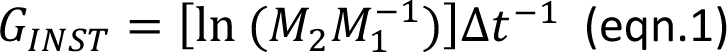

The change in fish’s total length (*TL_c_*) over time was measured using the total length at the beginning (*TL_1_*) and at the end (*TL_2_*) of the experimental period (*t*, number of days), following the equation 2:

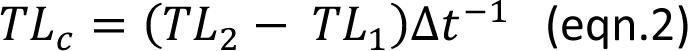

**(b) Foraging performance** was measured bite rate (total number of feeding strikes per minute). A siphon introduced approximately 1.5 grams of fish pellets to the test tank. The bite rate was recorded after removal of the siphon for 3 minutes. A clear action where the fish ingested the food pellet was interpreted as a feeding strike (O’Connor and Booth, 2021). If a fish did not feed during the observation time, the bite rate was equal to 0 and the time to feeding was recorded as 180 seconds.

**(c)** To test the **shelter response**, one cut-off section of a cylindrical PVC pipe was introduced into the test tank, and the shelter response was ranked based on different categories. 0 - Immediately fleeing away from structure, normal swimming patterns maintained. 1-No response to the structure, normal swimming patterns maintained. 2-No use of the structure until ≥60 sec, normal swimming behaviours modified to approach and enter the structure. 3-Immediate or almost immediate use of the structure <60 sec into the observation period, normal swimming behaviours modified to approach and enter the structure.

**(d)** A miniature block structure was introduced into the test tank with as minimal disturbance as possible to assess **fish boldness**. Block structures have been used in previous experiments to represent a novel object that does not emulate any natural structure that fish could have experienced in the wild (White *et al*., 2013). Fish boldness was ranked based on different categories. – 0 Immediately fleeing away from structure, swimming movements in opposite direction of the structure, no approach to the structure. -1 No response to the structure, normal swimming patterns maintained, no approach to the structure. -2 No investigation of the structure until ≥30 sec into the observation period, normal swimming behaviours modified to approach structure, mean approach distance 2-4 body lengths from structure. -3 Immediate or almost immediate investigation of the structure <30 sec into the observation period, closely approaches structure and initiates physical contact (i.e., feeding strikes or bumping), 1-2 body lengths from structure.

**(e)** A plastic fishing lure was used to mimic a predator to assess **predator escape response**. The fishing lure was dropped inside the test tank through a cable system to induce a flee response like the one that a predator would cause while minimising the potential effect of human presence (Djurichkovic *et al*., 2019; Figueira *et al*., 2019).

**(f) Burst-swimming speed** was calculated as the distance (cm) travelled in the 2 seconds following the fishing lure release. Fish lateralisation (left or right body turn) after disturbance was also assessed. Fish escape response was also ranked based on categories. -0 No response to disturbance, normal behaviour is maintained throughout the disturbance and afterwards, no change in swimming pattern. -1 Slight latency to disturbance (1-2 sec), initial escape response (<10 sec) is followed by an immediate return to normal behaviour, a moderate increase in swimming speed, erratic shifts in swimming angle. -2 Slight latency to disturbance (1-2 sec), initial escape response (<30 sec) is followed by an immediate return to normal behaviour, a moderate increase in swimming speed, erratic shifts in swimming angle. -3 Immediate or almost immediate response, escape response continues after disturbance (≥30 sec), large increase in swimming speed, substantial erratic behaviour, and/or followed by freezing behaviour.

**(g) Fish metabolism** is difficult to measure and as such, oxygen consumption rate is used as a proxy (McMahon *et al*., 2020). Resting metabolic rate (RMR) is evaluated by calculating the oxygen consumption of a fish while at rest (MO_2rest_), whereas the maximum metabolic rate (MMR) is estimated by calculating the maximum oxygen consumption (MO_2max_) when the fish is actively swimming. Aerobic scope was then calculated as the difference between the maximum and minimum oxygen consumption as per equation 3 from McMahon *et al*. (2020):

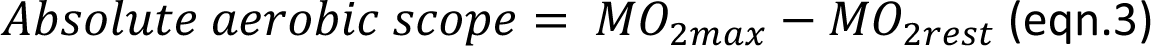

MO_2max_, MO_2rest_ and the absolute aerobic scope were calculated in mg O_2_ kg^-1^ h^-1^ following the equation 4 from McMahon *et al*. (2020):

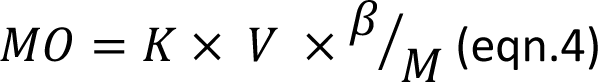

where *K* is the linear rate of decline (kPah^-1^) in the oxygen content over time (*h*) in the respirometry chamber, *V* is the volume of the chamber in L, 𝛽 is the water solubility (dependent on temperature and salinity, mg O_2_ L^-1^ kPa^-1^) and *M* is the mass of the fish (kg) (Munday *et al*., 2017).

Oxygen use in *G. subfasciatus*, and *P. sexlineatus* was measured as per Donelson et al. (2011) and McMahon et al. (2020) with intermittent respirometry.

**(h) Otolith analysis** was used to assess if fish growth in laboratory conditions reflected its recent growth performance in the wild before capture (Rigg *et al*., 2023). This aimed to assess if the laboratory growth results were representative and applicable to the field (Payne *et al*., 2016).

Otoliths were removed by dissection of the frontal bone as per Green *et al*. (2012) after fish were euthanised. Otoliths were then embedded in clear epoxy resin on a glass slide, placed anti-sulcus side up, and left to dry in the dark for at least 24 hours. Otoliths were polished to a plane using lapping films to reveal the peripheral growth increments.

A Nikon Eclipse *Ni* compound microscope was used to quantify otolith area, perimeter, diameter, and width of increments during the laboratory period and ten days before fish capture (in the wild).

### 2.5 Statistical analysis

To assess if there was a difference in performance between the two temperatures, and if there was an effect of order of temperature and/or food treatments, multifactorial ANOVAs were performed for each performance metric for each species. The first factor was the temperature, the second factor was the order of temperatures, and the experiments with *G. subfasciatus* and *P. sexlineatus* also included a third factor: food regimes. Interaction terms were also included.

To assess if there was a relationship between performance metrics, linear regressions were performed between pairs of continuous variables (instantaneous growth rate, change in total length, bite rate, burst speed and aerobic scope). Ordinal regressions were used when ordinal metrics were assessed (shelter response, boldness, escape response).

To assess if laboratory conditions provided equivalent fish growth conditions compared to the ones in the wild, otolith increment width (a proxy of fish growth) during laboratory conditions was compared to that over the 10 days prior to capture with paired t-tests. As water temperature at the time of *C. australis* collection was approximately 16°C, only the group of *C.australis* that was exposed to temperature in an ascending order was used for this analysis (increment width 10 days prior capture vs width at 16°C in the lab). Similarly, as water temperature at the time of *P. sexlineatus* collection was approximately 20°C, the group of *P. sexlineatus* that was exposed to temperature in a descending order was used (increment width 10 days prior capture vs width at 20°C in the lab). The assessment of *G. subfasciatus* otolith growth under laboratory conditions was omitted, primarily due to the discrepancy in water temperature at the time of collection (18°C), which did not align with any temperature treatment designated for the experiment. Given the acknowledged influence of temperature on otolith growth, maintaining consistent temperatures between field and laboratory environments would have been imperative to accurately determine any potential impact of the laboratory conditions on growth (Fey and Greszkiewicz, 2021). Assumptions of normality and homogeneity of variances were assessed with the Shapiro-Wilk normality test and Levene’s Equality of Error Variance test, respectively. Data that were found to violate the assumptions were log(x+1) or double-square root transformed, depending on requirements. Outliers were removed following the interquartile range method (Walfish, 2006).

The data were statistically analysed using SPSS Statistics (IMB Corporation, 2019) with a significance level (α) of 0.05.

## 3. Results

### 3.1 Fish population performance

Most performance metrics differed between the two test temperatures for *Gerres subfasciatus* and *Pelates sexlineatus*, but not *Centropogon australis* (Table 1).

**Table 1.**
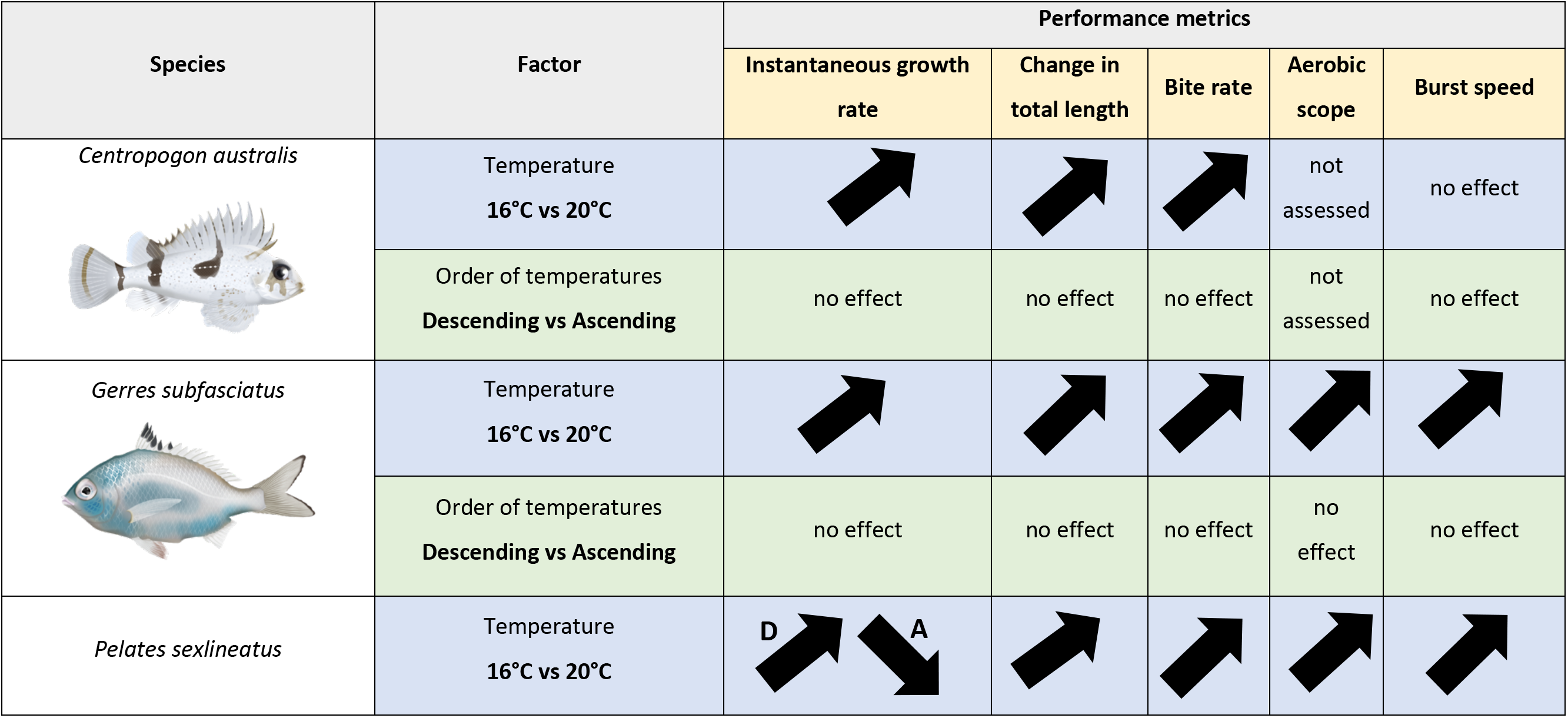
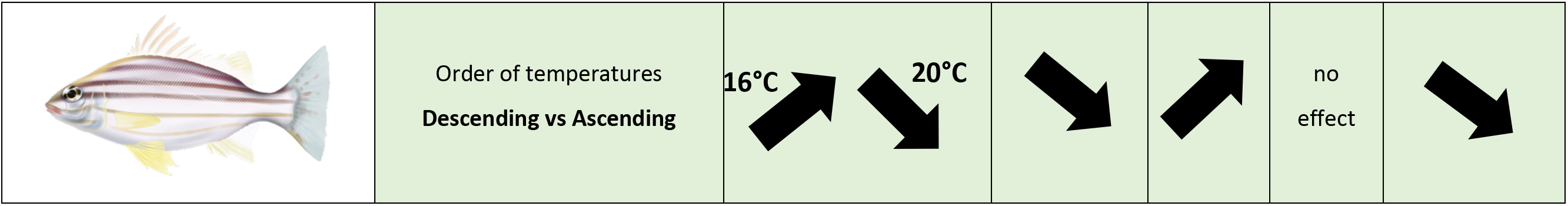
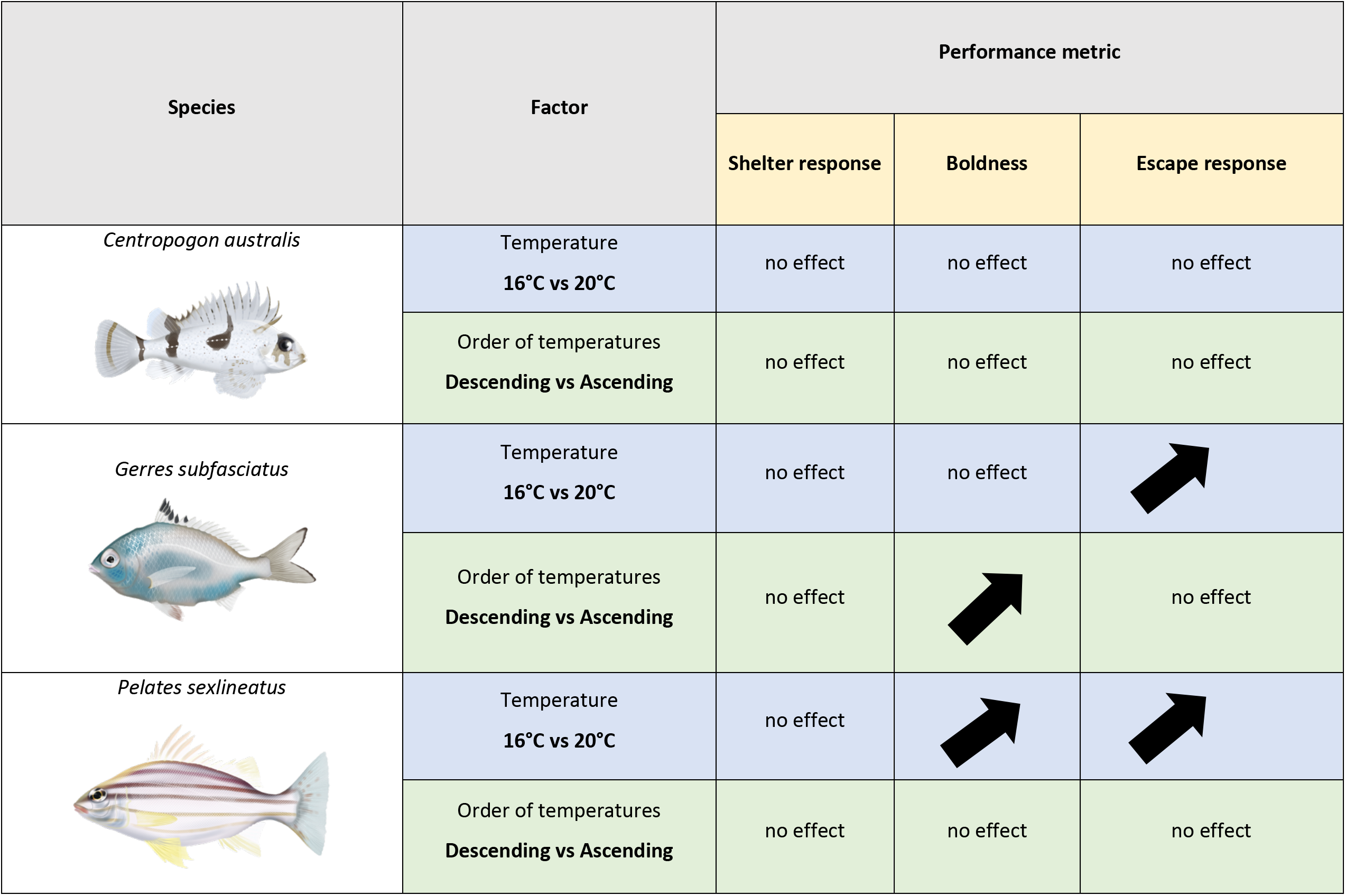
(continued at the top of following page). Summary of results obtained for *Centropogon australis, Gerres subfasciatus* and *Pelates sexlineatus* physiological and behavioural performance across treatments. Upward arrow indicates a significant increase in a performance metric with temperature/order of temperatures and refers downward arrow indicates a significant decrease in performance metric with temperature/ order of temperatures. **A** refers to Ascending temperature order (16°C to 20°C) and **D** refers to Descending temperature order (20°C to 16°C); these were used when the effect of temperature on the response variable significantly varied depending on the order of temperatures. The effect of food regimes was not significant and is discussed in the text. Please refer to supplementary materials (Figure S1 & Table S1) for detailed ANOVA outputs.

| Species | Factor | Performance metrics |  |  |  |  |
| --- | --- | --- | --- | --- | --- | --- |
|  |  | Instantaneous growth rate | Change in total length | Bite rate | Aerobic scope | Burst speed |
| *Centropogon australis* | Temperature<br>16°C vs 20°C | ↗ | ↗ | ↗ | not assessed | no effect |
|  | Order of temperatures<br>Descending vs Ascending | no effect | no effect | no effect | not assessed | no effect |
| *Gerres subfasciatus* | Temperature<br>16°C vs 20°C | ↗ | ↗ | ↗ | ↗ | ↗ |
|  | Order of temperatures<br>Descending vs Ascending | no effect | no effect | no effect | no effect | no effect |
| *Pelates sexlineatus* | Temperature<br>16°C vs 20°C | D ↗ A ↘ | ↗ | ↗ | ↗ | ↗ |
|  | Order of temperatures<br>**Descending vs Ascending** | 16°C ↗ 20°C ↘ | ↘ | ↗ | no<br>effect | ↘ |

| Species | Factor | Performance metric |  |  |
| --- | --- | --- | --- | --- |
|  |  | Shelter response | Boldness | Escape response |
| *Centropogon australis* | Temperature<br>16°C vs 20°C | no effect | no effect | no effect |
|  | Order of temperatures<br>Descending vs Ascending | no effect | no effect | no effect |
| *Gerres subfasciatus* | Temperature<br>16°C vs 20°C | no effect | no effect | ↗ |
|  | Order of temperatures<br>Descending vs Ascending | no effect | ↗ | no effect |
| *Pelates sexlineatus* | Temperature<br>16°C vs 20°C | no effect | ↗ | ↗ |
|  | Order of temperatures<br>Descending vs Ascending | no effect | no effect | no effect |

In terms of instantaneous growth rate, *C. australis* exhibited twice the growth rate at 20°C compared to 16°C. Similarly, for *G. subfasciatus,* growth was ten times higher at 20°C compared to 16°C. In *Pelates sexlineatus* an interaction was present between temperatures and order of temperatures, where the growth rate was higher at 20°C than at 16°C if fish were exposed to temperatures in a descending order (from 20°C to 16°C). In contrast, if fish were exposed to temperatures in an ascending order (16°C to 20°C), the growth rate was higher at 16°C than at 20°C. Therefore, the first temperature treatment to which *P. sexlineatus* were exposed post-capture presented the higher growth rates.

In *C. australis*, the change in total length was greater at 20°C compared to 16°C and in *G. subfasciatus*, the change in total length was seven times higher at 20°C than at 16°C. In *P. sexlineatus,* an interaction was present between temperatures and order of temperatures. The change in total length was always higher at 20°C than at 16°C, with the fish exposed to temperatures in an ascending order (from 16°C to 20°C) having a higher change in total length rates.

Aerobic scope was higher at 20°C than at 16°C in *G. subfasciatus* and in *P. sexlineatus*. Considering burst speed, this did not vary across treatments in *C. australis* while *G. subfasciatus* burst speed was higher at 20°C than at 16°C. In *P. sexlineatus,* an interaction was present between temperatures and order of temperatures. The burst speed was always higher at 20°C than at 16°C, with the fish exposed to temperatures in an ascending order (from 16°C to 20°C) having a higher burst speed.

The bite rate was nearly twice as high at 20°C compared to 16°C in both *C. australis* and *G. subfasciatus*. In *P. sexlineatus*, an interaction between temperatures and order of temperatures was present. Bite rate was always higher at 20°C than at 16°C with the fish exposed to temperatures in a descending order (from 20°C to 16°C) having higher bite rates. In *G. subfasciatus,* the order of administered temperatures influenced boldness which was higher in the descending temperature order treatments than in the ascending ones while escape response was higher at 20°C than at 16°C. In *P. sexlineatus* boldness and escape response were both higher at 20°C than at 16°C. While temperature and order of temperatures administered did not affect *C. australis* boldness or escape response. Across all species, no treatment affected fish shelter response.

### 3.2 Relationship between fish performance metrics

Most performance metrics were not related, particularly for *C*. *australis*, and significant relationships were generally weak (Table 2). In *C. australis*, only bite rate had a positive relationship with change in total length, while, across all other performance metrics the regressions were non-significant (0.07 < all *p-*values < 0.98). Since under 5% of the possible interactions were significant, these may have bene due to chance.

**Table 2.**
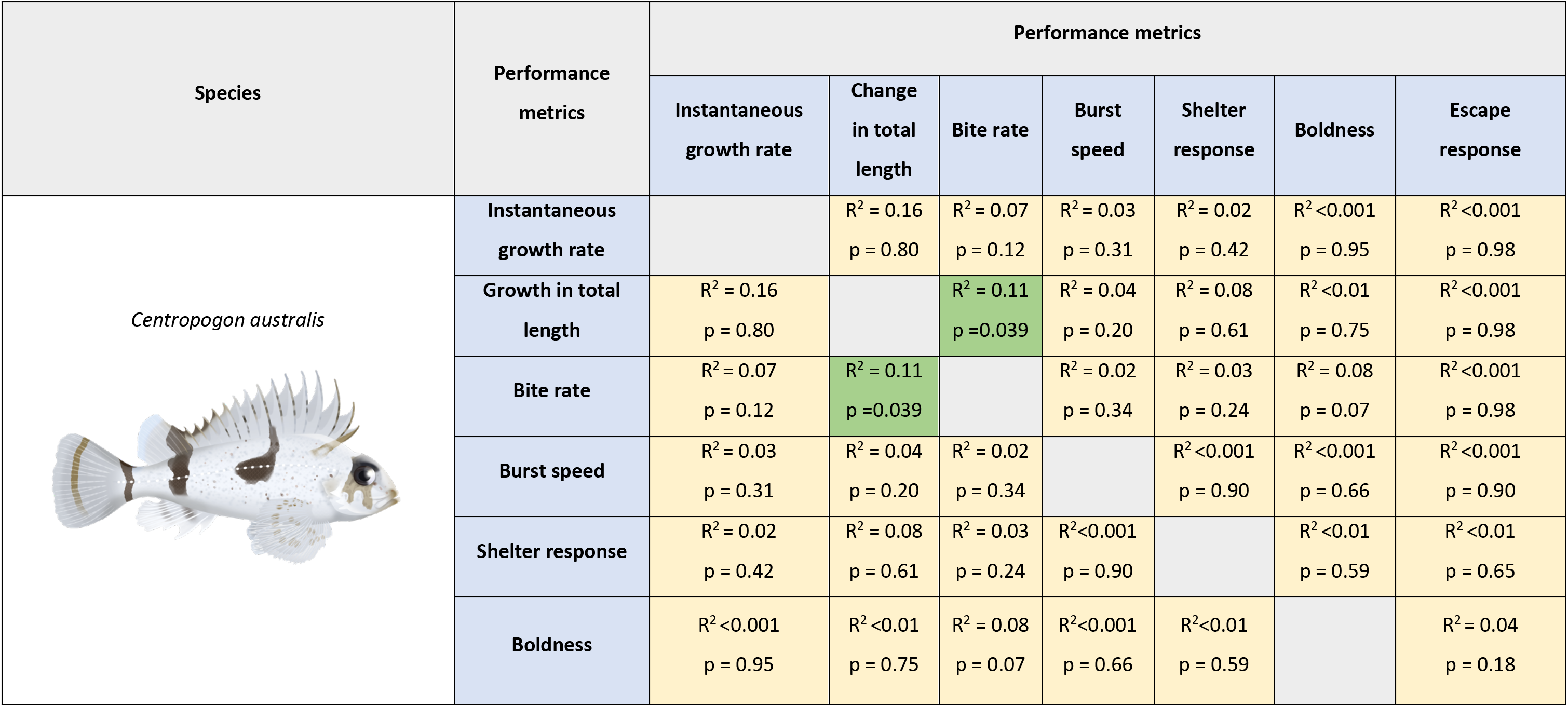
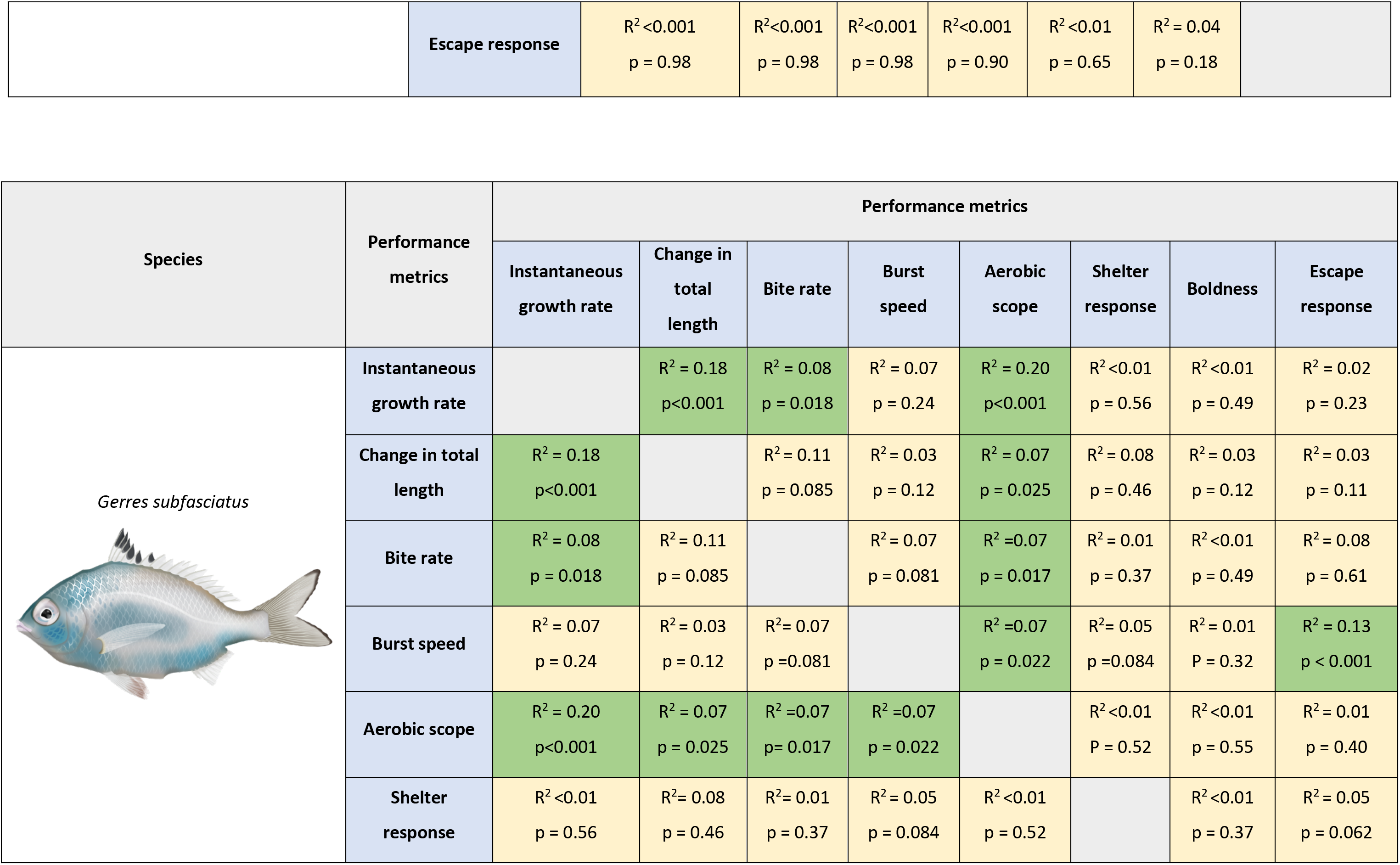
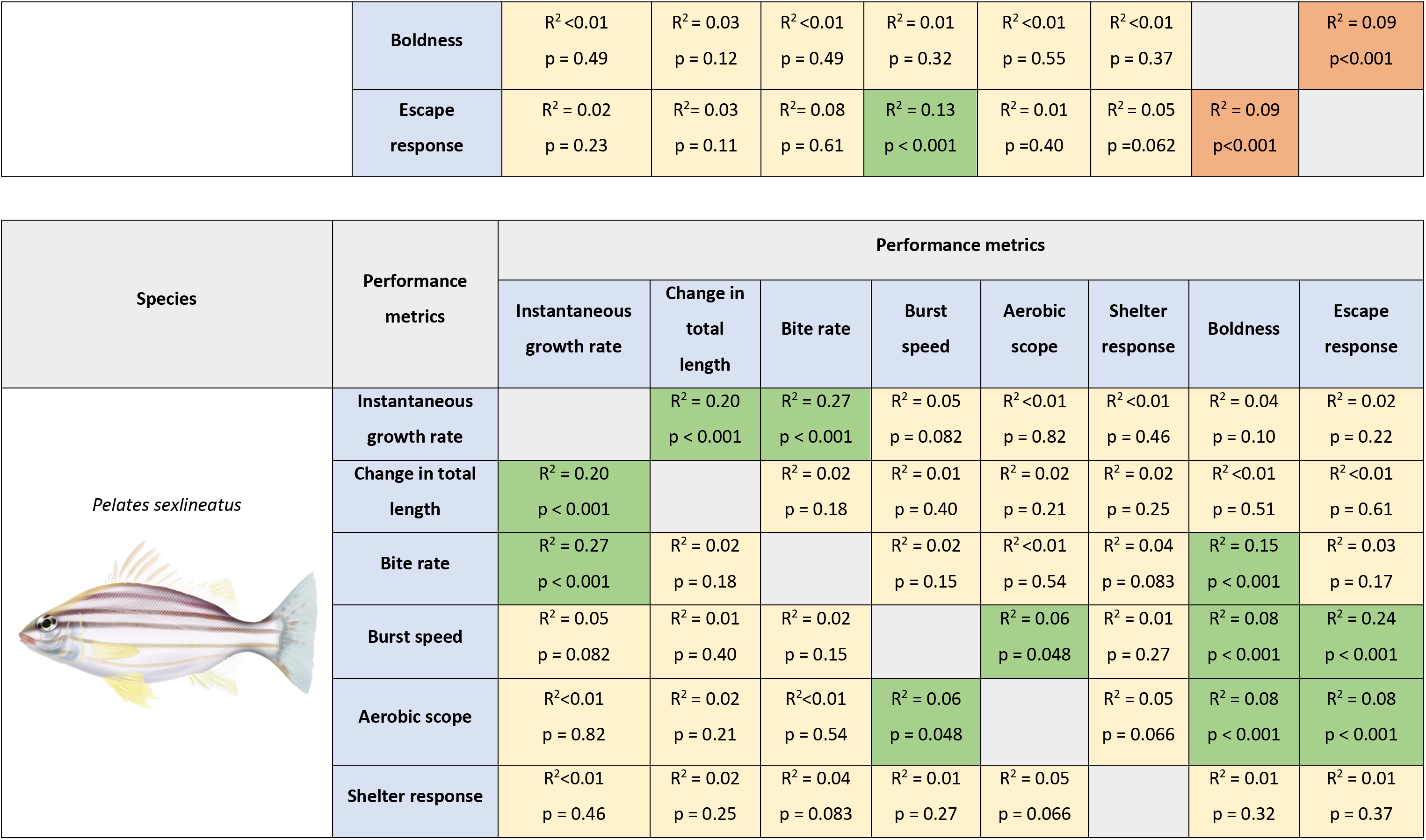
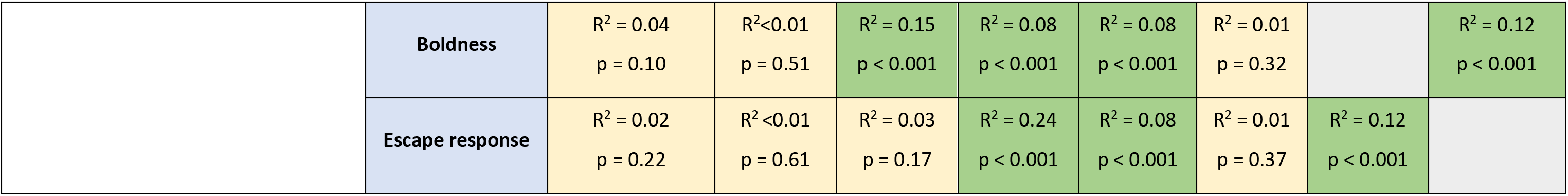
Results summary showing relationship patterns in *Centropogon australis*, *Gerres subfasciatus*, and *Pelates sexlineatu* physiological and behavioural performance regressions (performance metric 1 vs performance metric 2) with R^2^ and p-values. Red indicates a significantly negative relationship, green indicates a significantly positive relationship and yellow indicates the lack of a significant relationship. For linear regressions, Pearson’s R^2^ is reported, while for ordinal regressions, McFadden pseudo-R^2^ is reported. Please refer to supplementary materials (Table S2) for detailed ANOVA outputs.

| Species | Performance metrics | Performance metrics |  |  |  |  |  |  |
| --- | --- | --- | --- | --- | --- | --- | --- | --- |
|  |  | Instantaneous growth rate | Change in total length | Bite rate | Burst speed | Shelter response | Boldness | Escape response |
| *Centropogon australis* | Instantaneous growth rate | | $R^2 = 0.16$<br>$p = 0.80$ | $R^2 = 0.07$<br>$p = 0.12$ | $R^2 = 0.03$<br>$p = 0.31$ | $R^2 = 0.02$<br>$p = 0.42$ | $R^2 < 0.001$<br>$p = 0.95$ | $R^2 < 0.001$<br>$p = 0.98$ |
| | Growth in total length | $R^2 = 0.16$<br>$p = 0.80$ | | $R^2 = 0.11$<br>$p = 0.039$ | $R^2 = 0.04$<br>$p = 0.20$ | $R^2 = 0.08$<br>$p = 0.61$ | $R^2 < 0.01$<br>$p = 0.75$ | $R^2 < 0.001$<br>$p = 0.98$ |
| | Bite rate | $R^2 = 0.07$<br>$p = 0.12$ | $R^2 = 0.11$<br>$p = 0.039$ | | $R^2 = 0.02$<br>$p = 0.34$ | $R^2 = 0.03$<br>$p = 0.24$ | $R^2 = 0.08$<br>$p = 0.07$ | $R^2 < 0.001$<br>$p = 0.98$ |
| | Burst speed | $R^2 = 0.03$<br>$p = 0.31$ | $R^2 = 0.04$<br>$p = 0.20$ | $R^2 = 0.02$<br>$p = 0.34$ | | $R^2 < 0.001$<br>$p = 0.90$ | $R^2 < 0.001$<br>$p = 0.66$ | $R^2 < 0.001$<br>$p = 0.90$ |
| | Shelter response | $R^2 = 0.02$<br>$p = 0.42$ | $R^2 = 0.08$<br>$p = 0.61$ | $R^2 = 0.03$<br>$p = 0.24$ | $R^2 < 0.001$<br>$p = 0.90$ | | $R^2 < 0.01$<br>$p = 0.59$ | $R^2 < 0.01$<br>$p = 0.65$ |
| | Boldness | $R^2 < 0.001$<br>$p = 0.95$ | $R^2 < 0.01$<br>$p = 0.75$ | $R^2 = 0.08$<br>$p = 0.07$ | $R^2 < 0.001$<br>$p = 0.66$ | $R^2 < 0.01$<br>$p = 0.59$ | | $R^2 = 0.04$<br>$p = 0.18$ |
| | Escape response | $R^2 < 0.001$<br>$p = 0.98$ | $R^2 < 0.001$<br>$p = 0.98$ | $R^2 < 0.001$<br>$p = 0.98$ | $R^2 < 0.001$<br>$p = 0.90$ | $R^2 < 0.01$<br>$p = 0.65$ | $R^2 = 0.04$<br>$p = 0.18$ | |

| Species | Performance metrics | Performance metrics |  |  |  |  |  |  |  |
| --- | --- | --- | --- | --- | --- | --- | --- | --- | --- |
|  |  | Instantaneous growth rate | Change in total length | Bite rate | Burst speed | Aerobic scope | Shelter response | Boldness | Escape response |
| *Gerres subfasciatus* | Instantaneous growth rate | | $R^2 = 0.18$<br>$p < 0.001$ | $R^2 = 0.08$<br>$p = 0.018$ | $R^2 = 0.07$<br>$p = 0.24$ | $R^2 = 0.20$<br>$p < 0.001$ | $R^2 < 0.01$<br>$p = 0.56$ | $R^2 < 0.01$<br>$p = 0.49$ | $R^2 = 0.02$<br>$p = 0.23$ |
| | Change in total length | $R^2 = 0.18$<br>$p < 0.001$ | | $R^2 = 0.11$<br>$p = 0.085$ | $R^2 = 0.03$<br>$p = 0.12$ | $R^2 = 0.07$<br>$p = 0.025$ | $R^2 = 0.08$<br>$p = 0.46$ | $R^2 = 0.03$<br>$p = 0.12$ | $R^2 = 0.03$<br>$p = 0.11$ |
| | Bite rate | $R^2 = 0.08$<br>$p = 0.018$ | $R^2 = 0.11$<br>$p = 0.085$ | | $R^2 = 0.07$<br>$p = 0.081$ | $R^2 = 0.07$<br>$p = 0.017$ | $R^2 = 0.01$<br>$p = 0.37$ | $R^2 < 0.01$<br>$p = 0.49$ | $R^2 = 0.08$<br>$p = 0.61$ |
| | Burst speed | $R^2 = 0.07$<br>$p = 0.24$ | $R^2 = 0.03$<br>$p = 0.12$ | $R^2 = 0.07$<br>$p = 0.081$ | | $R^2 = 0.07$<br>$p = 0.022$ | $R^2 = 0.05$<br>$p = 0.084$ | $R^2 = 0.01$<br>$p = 0.32$ | $R^2 = 0.13$<br>$p < 0.001$ |
| | Aerobic scope | $R^2 = 0.20$<br>$p < 0.001$ | $R^2 = 0.07$<br>$p = 0.025$ | $R^2 = 0.07$<br>$p = 0.017$ | $R^2 = 0.07$<br>$p = 0.022$ | | $R^2 < 0.01$<br>$p = 0.52$ | $R^2 < 0.01$<br>$p = 0.55$ | $R^2 = 0.01$<br>$p = 0.40$ |
| | Shelter response | $R^2 < 0.01$<br>$p = 0.56$ | $R^2 = 0.08$<br>$p = 0.46$ | $R^2 = 0.01$<br>$p = 0.37$ | $R^2 = 0.05$<br>$p = 0.084$ | $R^2 < 0.01$<br>$p = 0.52$ | | $R^2 < 0.01$<br>$p = 0.37$ | $R^2 = 0.05$<br>$p = 0.062$ |
| | **Boldness** | $R^2 < 0.01$<br>$p = 0.49$ | $R^2 = 0.03$<br>$p = 0.12$ | $R^2 < 0.01$<br>$p = 0.49$ | $R^2 = 0.01$<br>$p = 0.32$ | $R^2 < 0.01$<br>$p = 0.55$ | $R^2 < 0.01$<br>$p = 0.37$ | | $R^2 = 0.09$<br>$p < 0.001$ |
| | **Escape response** | $R^2 = 0.02$<br>$p = 0.23$ | $R^2 = 0.03$<br>$p = 0.11$ | $R^2 = 0.08$<br>$p = 0.61$ | $R^2 = 0.13$<br>$p < 0.001$ | $R^2 = 0.01$<br>$p = 0.40$ | $R^2 = 0.05$<br>$p = 0.062$ | $R^2 = 0.09$<br>$p < 0.001$ | |
|  |  | Instantaneous growth rate | Change in total length | Bite rate | Burst speed | Aerobic scope | Shelter response | Boldness | Escape response |
| *Pelates sexlineatus* | Instantaneous growth rate | | $R^2 = 0.20$<br>$p < 0.001$ | $R^2 = 0.27$<br>$p < 0.001$ | $R^2 = 0.05$<br>$p = 0.082$ | $R^2 < 0.01$<br>$p = 0.82$ | $R^2 < 0.01$<br>$p = 0.46$ | $R^2 = 0.04$<br>$p = 0.10$ | $R^2 = 0.02$<br>$p = 0.22$ |
| | Change in total length | $R^2 = 0.20$<br>$p < 0.001$ | | $R^2 = 0.02$<br>$p = 0.18$ | $R^2 = 0.01$<br>$p = 0.40$ | $R^2 = 0.02$<br>$p = 0.21$ | $R^2 = 0.02$<br>$p = 0.25$ | $R^2 < 0.01$<br>$p = 0.51$ | $R^2 < 0.01$<br>$p = 0.61$ |
| | Bite rate | $R^2 = 0.27$<br>$p < 0.001$ | $R^2 = 0.02$<br>$p = 0.18$ | | $R^2 = 0.02$<br>$p = 0.15$ | $R^2 < 0.01$<br>$p = 0.54$ | $R^2 = 0.04$<br>$p = 0.083$ | $R^2 = 0.15$<br>$p < 0.001$ | $R^2 = 0.03$<br>$p = 0.17$ |
| | Burst speed | $R^2 = 0.05$<br>$p = 0.082$ | $R^2 = 0.01$<br>$p = 0.40$ | $R^2 = 0.02$<br>$p = 0.15$ | | $R^2 = 0.06$<br>$p = 0.048$ | $R^2 = 0.01$<br>$p = 0.27$ | $R^2 = 0.08$<br>$p < 0.001$ | $R^2 = 0.24$<br>$p < 0.001$ |
| | Aerobic scope | $R^2 < 0.01$<br>$p = 0.82$ | $R^2 = 0.02$<br>$p = 0.21$ | $R^2 < 0.01$<br>$p = 0.54$ | $R^2 = 0.06$<br>$p = 0.048$ | | $R^2 = 0.05$<br>$p = 0.066$ | $R^2 = 0.08$<br>$p < 0.001$ | $R^2 = 0.08$<br>$p < 0.001$ |
| | Shelter response | $R^2 < 0.01$<br>$p = 0.46$ | $R^2 = 0.02$<br>$p = 0.25$ | $R^2 = 0.04$<br>$p = 0.083$ | $R^2 = 0.01$<br>$p = 0.27$ | $R^2 = 0.05$<br>$p = 0.066$ | | $R^2 = 0.01$<br>$p = 0.32$ | $R^2 = 0.01$<br>$p = 0.37$ |
| | **Boldness** | $R^2 = 0.04$<br>$p = 0.10$ | $R^2 < 0.01$<br>$p = 0.51$ | $R^2 = 0.15$<br>$p < 0.001$ | $R^2 = 0.08$<br>$p < 0.001$ | $R^2 = 0.08$<br>$p < 0.001$ | $R^2 = 0.01$<br>$p = 0.32$ | | $R^2 = 0.12$<br>$p < 0.001$ |
| | **Escape response** | $R^2 = 0.02$<br>$p = 0.22$ | $R^2 < 0.01$<br>$p = 0.61$ | $R^2 = 0.03$<br>$p = 0.17$ | $R^2 = 0.24$<br>$p < 0.001$ | $R^2 = 0.08$<br>$p < 0.001$ | $R^2 = 0.01$<br>$p = 0.37$ | $R^2 = 0.12$<br>$p < 0.001$ | |

In *G. subfasciatus,* aerobic scope was positively related to instantaneous growth rate, change in total length, bite rate, and burst speed, while escape response was positively related to burst speed and negatively correlated with boldness. Additionally, instantaneous growth rate was positively related to change in total length and bite rate.

In *P. sexlineatus,* aerobic scope was positively related to burst speed, boldness, and escape response; burst speed was positively related to boldness and escape response; bite rate and escape response were positively related to boldness and instantaneous growth rate was positively related to bite rate and change in total length.

Fish growth under laboratory conditions was shown to be similar to fish growth in the wild with an otolith growth analysis (see Supplementary materials, Figure S3).

## 4. Discussion

Overall, as expected from metabolic theory, performance was generally higher at the higher water temperature for all fish species, and this suggests that winter performance of juveniles for these temperate estuarine species may be enhanced under future warming. Food regimes did not have an effect on fish performance, suggesting temperature responses may not be mediated by food availability. There was also no strong relationship among metrics, which was occasionally counter-intuitive given expected relationships (e.g. expected negative relationship between boldness and sheltering), suggesting physiological and behavioural responses to warming may be complex and species-specific.

Generally, as temperature increases without exceeding the thermal optimum, metabolic rate increases, and behavioural and physiological processes are stimulated, inducing higher growth rate, development times and swimming activity (Kent and Ojanguren, 2015; Tsoukali *et al*., 2016). The aerobic scope generally follows the bell-shape performance curve pattern, increasing at moderate temperatures toward the thermally optimal temperature but then declining (Norin *et al*., 2014). A reduction in growth rates and aerobic scope with cooler temperatures, along with a decrease in activity, is a strategy to allow individuals to increase the surplus of aerobic scope that can be directed towards coping with environmental stressors (e.g., increase metabolic heat production) (Killen, 2014). As expected, *Gerres subfasciatus* and *Pelates sexlineatus* presented higher aerobic scope at 20°C than at 16°C and *G. subfasciatus* presented higher growth rates at 20°C. This suggests that our experimental temperatures were below the thermal optimum for these two species (Paloheimo and Dickie, 1966; Reist *et al*., 2006; Thresher *et al*., 2007; Neuheimer *et al*., 2011; O’Connor and Booth, 2021). This was expected as the tested temperatures are within the species thermal range (Booth et al. 2014).

Additionally, a positive relationship was detected between aerobic scope and growth rates in *G.subfasciatus.* Higher aerobic scope potentially afforded more energy above basal metabolic requirements for growth in this species. This might be expected since individuals within their optimal temperature range and with higher metabolic rates should also have higher scope for growth (Killen *et al*., 2011). Similar results were found by Auer *et al*. (2015), McCarthy (2000) and Álvarez and Nicieza (2005) where higher standard metabolic rates were associated with faster growth rates in juvenile *S. trutta, S salar,* and *S.trutta*. Surprisingly, no relationship was detected between aerobic scope and growth rates in *P. sexlineatus*. This might indicate that the energy surplus derived from higher aerobic scope was diverted to other processes such as activity and responsiveness to stimuli (Paschke *et al*., 2018), however, this species was tested at temperatures well below its thermal optimum where performance is maximised (Booth *et al*., 2014).

A slight rise in winter water temperature in the study region may therefore be beneficial for the growth performance of juveniles of *G. subfasciatus*, *P. sexlineatus* and *Centropogon australis* which increased with temperature. Previous studies that assessed *P. sexlineatus* and *C. australis* growth at 18°C, 22°C and 26°C found that growth was highest at 22°C and 26°C respectively (Booth *et al*., 2014), thus our tested temperatures were below their thermal maxima. Providing that the temperature remains below the thermal optimum for each species, future temperature increases in the region are likely to increase energy availability, result in faster rates of diffusion, and more enzyme-substrate complexes, which will ultimately lead to higher reaction rates and increased somatic growth (Neuheimer *et al*., 2011). If the increased growth rate continues into later developmental stages, it may reduce size and age at maturation and resulting recruitment dynamics (Kuparinen *et al*., 2011).

The lower growth rates at cooler temperatures could be explained by the increased mitochondrial oxidative capacities and mitochondrial proliferation (Guderley, 2004; Kammer *et al*., 2011). For instance, the cold-acclimated three-spined stickleback (*Gasterosteus aculeatus*) had a higher protein damage induced by ROS (reactive oxygen species) proliferation than the warm-acclimated group (Kammer *et al*., 2011). An increase in ROS production and oxidative damage can decrease total animal performance. For example, by reducing swimming performance and muscle function (Ghanizadeh Kazerouni *et al*., 2016). This could explain the decrease in burst speed observed in *G.subfasciatus* and *P. sexlineatus* in the lower temperature treatment.

The high energetic costs required for biosynthetic processes that allow for growth imply that behavioural patterns enhancing food intake rates should positively relate to growth (Biro and Stamps, 2010). Foraging activity increased with temperature across all species in the current study, which was likely motivated by the higher metabolic requirements that come with higher temperatures (Ferreira *et al*., 1998; Byström *et al*., 2006; Smith, 2008; Nowicki *et al*., 2012; O’Connor and Booth, 2021). Fish use sensory systems like hearing, olfaction/taste, vision, and cognitive abilities for food detection. Appetite-regulating endocrine factors, released by organs influencing brain feeding centres, control food intake (Volkoff and Rønnestad, 2020). Temperature changes affect these pathways, altering feeding behaviour and taste preferences. Cold water reduces nutrient digestibility by modifying digestive enzyme activity, typically increasing digestion rates and gut transit time while decreasing gastrointestinal evacuation rates (Volkoff and Rønnestad, 2020). This may explain lower bite rates at lower temperatures and a positive correlation between bite rate and growth across species.

A change in growth (mass) is usually accompanied by a change in length (Jones et al., 1999), and it was observed that higher growth rates at higher temperatures were accompanied by greater increases in total length in *P.sexlineatus* individuals exposed to temperatures in a descending order, and in *G.subfasciatus* irrespective of order of exposure. While for *C.australis*, an increase in length with temperature was decoupled from an increase in mass, indicating that the change in temperature might have induced a modification in the body shape (e.g., depth, elongation) that did not strongly affect body mass (Poléo *et al*., 1995). Such morphological modifications can induce differences in behaviours as predator evasion, foraging behaviour, activity level and explorativeness (Ahti *et al*., 2020).

We found that lab growth performance reflected recent field growth prior to collection for *P. sexlineatus* and *C. australis*, suggesting that laboratory artefacts on growth rate were minimal in the current study. This has been found in other studies that assessed fish DNA (Fukuda *et al*., 2001), and aerobic scope (Payne *et al*., 2016), with individual laboratory growth sufficiently representing growth in the wild. While these findings together suggest that tank experiments may be suitable for growth-related experiments for fishes, we were unable to test other confounding effects potentially arising from laboratory conditions, e.g. behavioural modification. Given the potential for environmental factors and ecological interactions (e.g. predation) to mediate behavioural responses (Beckerman *et al*., 2007), we therefore cannot eliminate the possibility of potential biases for behavioural metrics in the current study.

A positive relationship between aerobic scope and burst speed was found in *P. sexlineatus*; this could be explained by the fact that the surplus of energy that comes with a higher aerobic scope can be invested in energetically expensive tissues that boost locomotor performance (Metcalfe *et al*., 2016). Higher aerobic scope increases the fish capacity for aerobic swimming performance with higher critical swimming speeds, as seen in the cod *Gadus morhua* (Reidy *et al*., 2000) and aerobic scope was positively correlated with gait transition speed in the mullet *Liza aurata* (Killen *et al*., 2012). In *P. sexlineatus*, growth rate decreased in the second temperature treatment irrespective of the temperature. Therefore, the lower performance might also be due to an effect of laboratory conditions on the specimens rather than solely temperature. The capture, handling and change of environment can induce psychological and physical stress that can impact fish growth and survival (Brydges *et al*., 2009). However, the fish were monitored throughout the experiment and appeared healthy, behaving in a non-stressed manner. Additionally, in *P. sexlineatus* individuals exposed to temperatures in an ascending order, the change in total length and burst speed were higher while bite rate was lower compared to the fish exposed to temperatures in a descending order. The initial exposure to cold temperatures might induce long-lasting adverse effects on fish physiological processes (Donaldson *et al*., 2008). This effect was not observed for the other two species in the current study, indicating potentially complex and species-specific responses to temperature change.

In accordance with our results on *G. subfasciatus*, other studies have found a positive relationship between foraging activity and aerobic scope, as individuals with higher metabolic rates had higher foraging requirements to support the increased energy demand (Killen, 2014). Metabolism has been previously shown to influence behavioural traits such as dominance, aggression and boldness in fish (Metcalfe *et al*., 2016). In accordance with our results on *P. sexlineatus,* previous studies have found a positive association between aerobic scope and boldness. For instance, the *S. salar* fry that moved further away from the nest had higher metabolic rates (measured during the egg stage) (Robertsen *et al*., 2014), while the common carp *Cyprinus carpio* individuals that presented the highest resting metabolic rates also had the riskier behaviour (Huntingford *et al*., 2010).

In accordance with our results for *P. sexlineatus*, various studies with other species, such as the damselfish *Pomacentrus moluccensis*, the Amazonia cichlid *Apistogramma agassizii,* and the Siamese fighting fish *Betta splendens*, revealed that fish presented higher boldness and activity rates with an increase in temperature (Clarke and Johnston, 1999; Hölker, 2006; Biro *et al*., 2010; Kochhann *et al*., 2015; Forsatkar *et al*., 2016; Laubenstein *et al*., 2018). Warmer temperatures modify energy budgets and risk assessment in fish, increasing the activity levels of individuals while enhancing their predation risk (Forsatkar *et al*., 2016). Thus, bolder fish are less likely to have a prompt response to escape predators. Higher mortality rates in the rainbow trout, *O. mykiss* across lakes in Canada was attributed to the increase in temperature that boosted fish activity levels and risk-taking behaviours compared to cooler years (Biro *et al*., 2007). This was explained by the fact that fish are compensating for the increased metabolic cost that is associated with increased temperature and thus require higher activity rates to increase the food intakes that are necessary to maintain the preferred (intrinsic) growth rates at the higher temperatures (Biro and Stamps, 2008).

Surprisingly, the order of temperature treatment affected *G. subfasciatus* boldness, with the fish being exposed to the warmer (20°C) treatment first being bolder than the ones that experienced 20°C temperatures second (ascending order). The negative effect of colder temperature on boldness experienced by the fish in the ascending order group might have persisted when they were exposed to the second (warmer) temperature rendering them less active and resilient than the other group (Gingerich *et al*., 2007; Kim *et al*., 2019).

Skeletal muscle, specifically, displays robust adaptations that frequently result in temperature-related compensation of movement ability with an increase in force production, relaxation, shortening and production of power output with initial increases in temperature (James and Tallis, 2019). Various studies have described an increase in escape response and swimming speeds at higher temperatures (Grigaltchik *et al*., 2012; Djurichkovic *et al*., 2019; Domenici and Hale, 2019). These are consistent with our findings with *P. sexlineatus* and *G. subfasciatus* which indicate higher performance at higher temperatures whilst no changes with temperatures were observed in *C. australis*. Temperature affects swimming propulsive performance as the muscle power output increases with higher temperatures (Wilson *et al*., 2010). Fast starts rely on anaerobic power, but the energy debt incurred during this process must be compensated through post-exercise oxygen consumption. This process requires more energy than was initially used, resulting in an energy deficit (Allan *et al*., 2015). Thus, the higher aerobic scope that is present at higher temperatures can provide a surplus of energy that can be invested into greater performance (in our case, as both boldness and escape response) (Bergman, 1987).

Furthermore, temperature influences the fish’s neural processing times and thus affects escape latency and directionality (Domenici and Hale, 2019). Colder temperatures are associated with increased response latency and lower peak acceleration and velocity due to a reduction in initial fast body bend away from the stimulus (predator), thus escape response (Beitinger *et al*., 2000). At a cellular level, this can be explained by the fact that cooling induces an increase in the Mauthner neurons input resistance (a brainstem network that triggers escape response), causing alterations in the balance between the inhibitory and excitatory transmission onto the Mauthner neurons cell system (Preuss and Faber, 2003; Szabo *et al*., 2008).

Some studies have highlighted how escape response changes if the fish are part of a school, which usually slows escape response, likely due to the low perceived risk of predation in a group (Domenici and Hale, 2019). As my study placed fish in individual confinement to assess individual performance, future studies could group fish together to assess density-dependent effects on these performance functions. Group-size effects might be particularly relevant for social shoaling species (*P. sexlineatus* and *G. subfasciatus*) (Sbragaglia *et al*., 2021).

As expected, a negative relationship between boldness and escape response was detected in *G. subfasciatus,* and this would indicate that the bolder individuals also have lower escape response and thus might be more likely to be predated upon than less bold individuals (Killen *et al*., 2011) at winter temperatures. This might represent a trait compensation situation whereby the costs of one trait are offset by the benefits of another trait (Kuo *et al*., 2015). In this instance, the higher foraging success and mate encounter chances that come with high boldness are counteracted by the higher vulnerability to predators that comes with a lower escape response (Paijmans *et al*., 2020). In comparison, *P.sexlineatus* had a positive relationship between boldness and predator escape response indicating that the higher activity and risk-taking behaviour that comes with boldness was coupled with higher responsiveness to stimuli such as predators (Ioannou *et al*., 2008). For *C. australis*, no relationship was found across these metrics. The variations in responses among species hint at potential intricate and species-specific reactions to temperature fluctuations.

The ability to escape predators and thus have higher survival rates is affected not only by a fish’s responsiveness and burst speed but also by the strategic use of habitat resources, such as shelter (Halpin, 2000). Shelter is a vital resource that can influence survival through evading predators. Shelter response did not significantly differ across treatments for any species. However, trade-offs exist between predator avoidance and foraging or mating success (Finstad *et al*., 2007). Some individuals might increase their risk of starvation if the shelter usage limits the time spent foraging, while others might increase their foraging efficiency by sheltering and ambushing preys (Conallin *et al*., 2012). The latter could explain the ambush predator *C.australis* relatively low boldness and high shelter usage compared to the other species (Bell *et al*., 1978).

As trophic interactions rely on the relative body size of predators and prey, even slight variations in how species respond to changes in ambient temperatures could alter the structure and function of marine systems by inducing a mismatch in prey and predator phenotype with potentially adverse effects on predator-prey dynamics and thus trophic interactions (Grigaltchik *et al*., 2012; Durant *et al*., 2019). When species at different trophic levels exhibit different growth rates, it can change the energetic requirements, prey preferences, and top-down regulation of apex or mesopredators (Van Denderen *et al*., 2020). Additionally, the vulnerability of certain species to predation may also be affected, leading to a chain reaction of consequences throughout the entire ecosystem (Pistevos *et al*., 2015). Changes in the organization of marine systems might be amplified if the foraging capabilities of fish species are also impacted by temperature, as even minor alterations to this behaviour can significantly impact survival. It is widely recognized that fish reared in warmer temperatures tend to exhibit higher levels of aggression, feeding rates, foraging activity, and escape responses (Domenici, 2010). Thus these changes in behaviour will likely have strong effects on future community dynamics, and future studies could implement multi-species assessments (Domenici and Hale, 2019). The disparities observed in performance responses across species, notably the contrast between *C. australis* and *G. subfasciatus* and *P. sexlineatus*, could underscore the potential differential impacts of climate change on estuarine fish communities in the future. This variability in responses among species may exert significant influences on the overall ecosystem’s trophic interactions and dynamics.

Overall, we found that temperature had a pervasive effect (more than food regime or order of temperatures) on fish performance, thus thermal physiology (Little *et al*., 2020). We did not observe significant interactions between temperature and food. This suggests that, contrary to expectations (Donelson *et al*., 2010; McLeod *et al*., 2013), performance limitations due to food scarcity did not occur, at least within the parameters of our experimental conditions. Fish generally performing better at higher temperatures which might indicate that the tested temperatures were below the species’ thermal optima. In light of climate change-induced sea temperature rise, the wide thermo-tolerance and physiological adaptability of these estuarine species that enables them to inhabit such dynamic systems might be instrumental in sustaining the health and productivity of estuaries and supporting the fisheries that depend on them (Booth *et al*., 2014). However, differences between species were present; this suggests that juvenile stages of estuarine species will react differently to climate change. Thus, the overall effects on ecosystem dynamics and productivity are difficult to predict (O’Connor and Booth, 2021). To further improve the knowledge of estuarine community response to climate change, future studies could encompass other estuarine species (Astles and McLeod, 2018) and multi-species/predator-prey interactions (Nagelkerken *et al*., 2016).

## Acknowledgments

We would like to thank the University of Technology, Sydney (UTS) for supporting and providing the facilities for this project. We would like to firstly thank Dr Jennifer Donelson, who provided essential insights in how to measure fish metabolism and Prof William Figueira who generously lent us the oxygen-measuring equipment. Special thanks to the laboratory technical staff Helen Price, Rachel Keeys, Scott Allchin, and Sue Fenech for promptly assisting me with any laboratory issue.

## Supporting information

Supporting information available as attached file.

## Contributions

All authors contributed to data generation, analysis, manuscript preparation, and conceptualizing the paper’s structure.

