## Supplementary materials for "Overwintering performance of juvenile temperate estuarine fish"

**<u>Bellotto et al Supplementary data</u>**

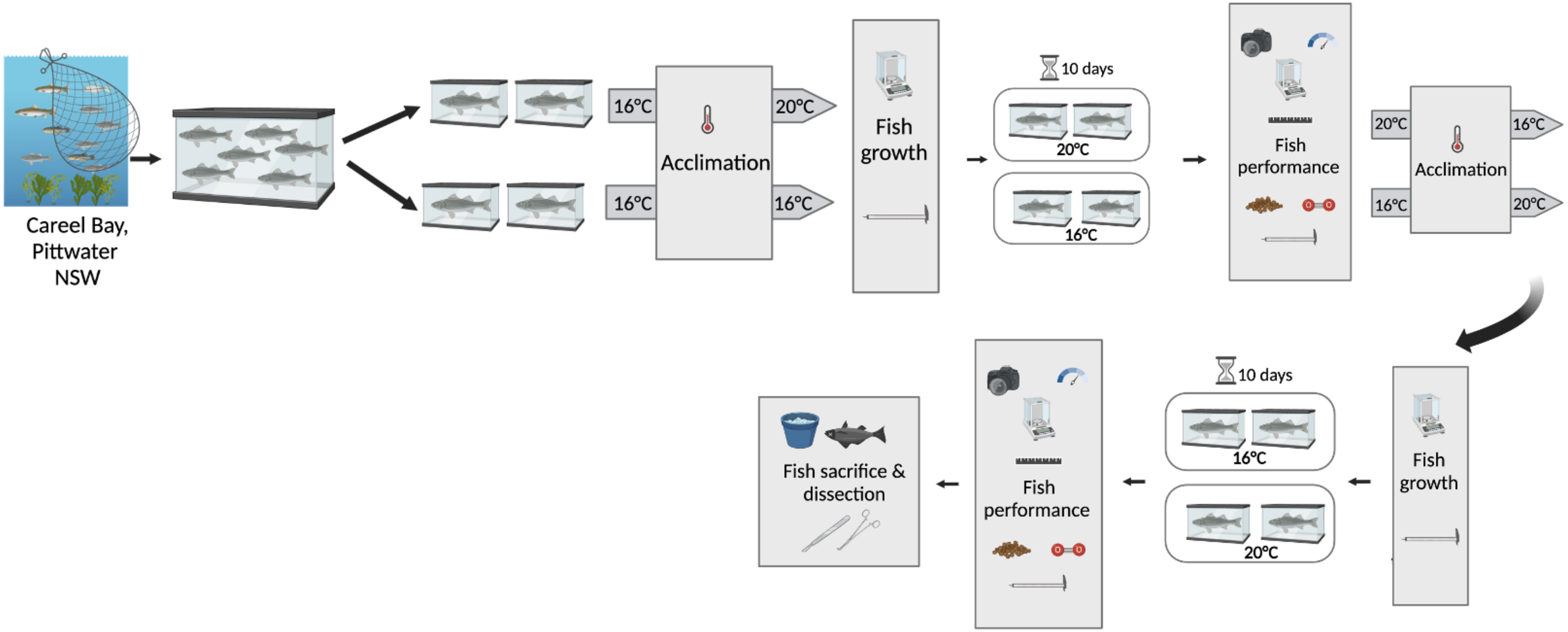


**(A)**

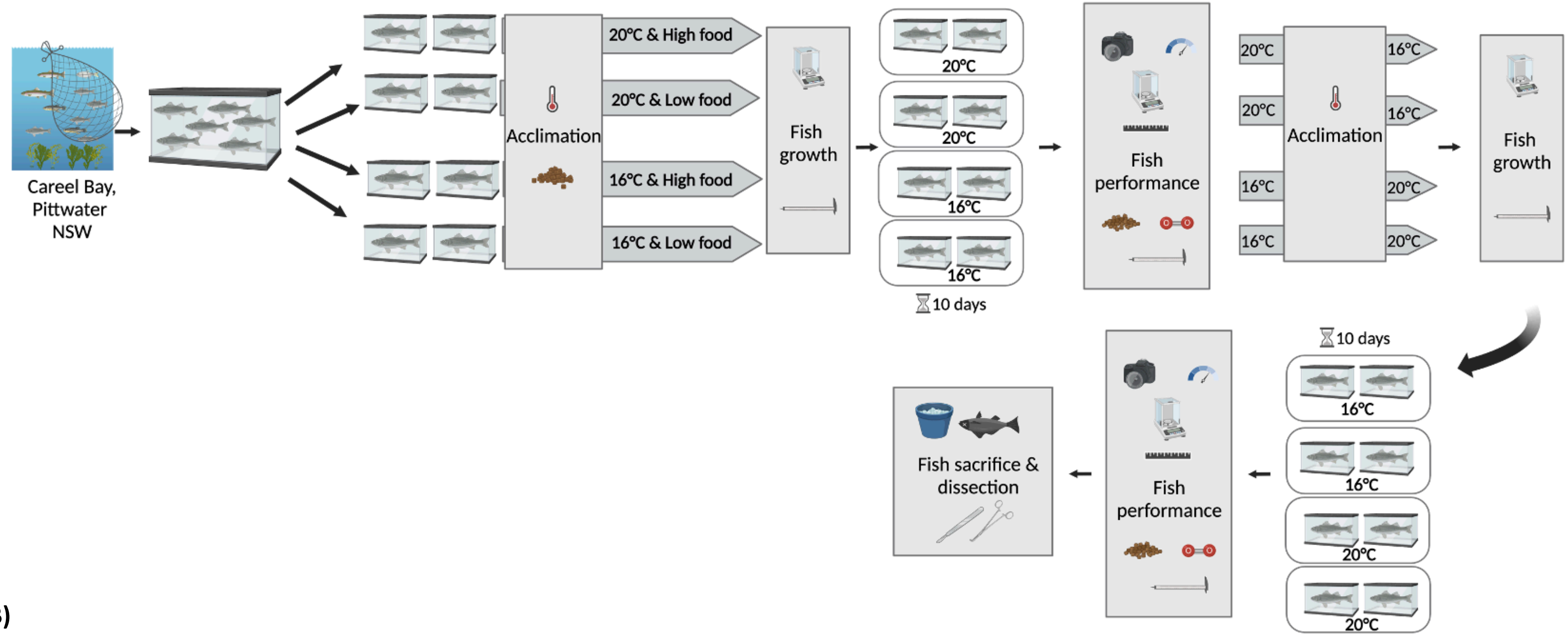


**(B)**

**Figure S1. (A)** Diagram of experimental design for the eastern fortescues, *Centropogon australis* (n = 20), with temperature (16°C vs 20°C) as factor. **(B)** Diagram of experimental design for the common silver biddy *Gerres subfasciatus* (n=37) and the eastern striped trumpeter, *Pelates sexlineatus* (n=34) with temperature (16°C vs 20°C) and food (low vs high food regime) as factors. A separate experiment was conducted for each species. Low food regimes were fed 0.5% fish weight in fish pellets, High food regimes were fed 1% fish weight in fish pellets. Created with BioRender (2023).

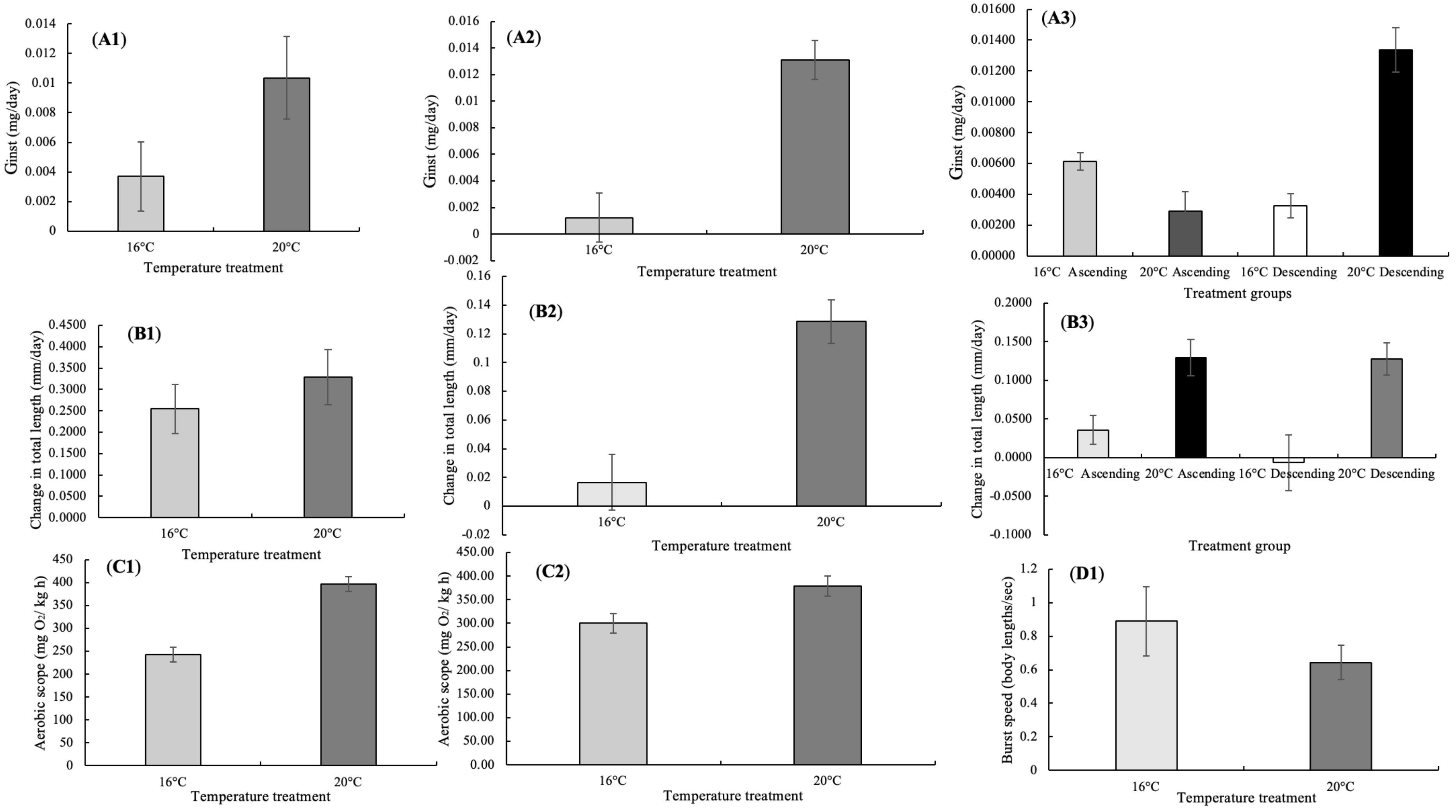

(A1)
Ginst (mg/day)
0.014
0.012
0.01
0.008
0.006
0.004
0.002
0
16°C
20°C
Temperature treatment
(A2)
Ginst (mg/day)
0.016
0.014
0.012
0.01
0.008
0.006
0.004
0.002
0
-0.002
16°C
20°C
Temperature treatment
(A3)
Ginst (mg/day)
0.01600
0.01400
0.01200
0.01000
0.00800
0.00600
0.00400
0.00200
0.00000
16°C Ascending
20°C Ascending
16°C Descending
20°C Descending
Treatment groups
(B1)
Change in total length (mm/day)
0.4500
0.4000
0.3500
0.3000
0.2500
0.2000
0.1500
0.1000
0.0500
0.0000
16°C
20°C
Temperature treatment
(B2)
Change in total length (mm/day)
0.16
0.14
0.12
0.1
0.08
0.06
0.04
0.02
0
-0.02
16°C
20°C
Temperature treatment
(B3)
Change in total length (mm/day)
0.2000
0.1500
0.1000
0.0500
0.0000
-0.0500
-0.1000
16°C Ascending
20°C Ascending
16°C Descending
20°C Descending
Treatment group
(C1)
Aerobic scope (mg O2/ kg h)
450
400
350
300
250
200
150
100
50
0
16°C
20°C
Temperature treatment
(C2)
Aerobic scope (mg O2/ kg h)
450.00
400.00
350.00
300.00
250.00
200.00
150.00
100.00
50.00
0.00
16°C
20°C
Temperature treatment
(D1)
Burst speed (body lengths/sec)
1.2
1
0.8
0.6
0.4
0.2
0
16°C
20°C
Temperature treatment

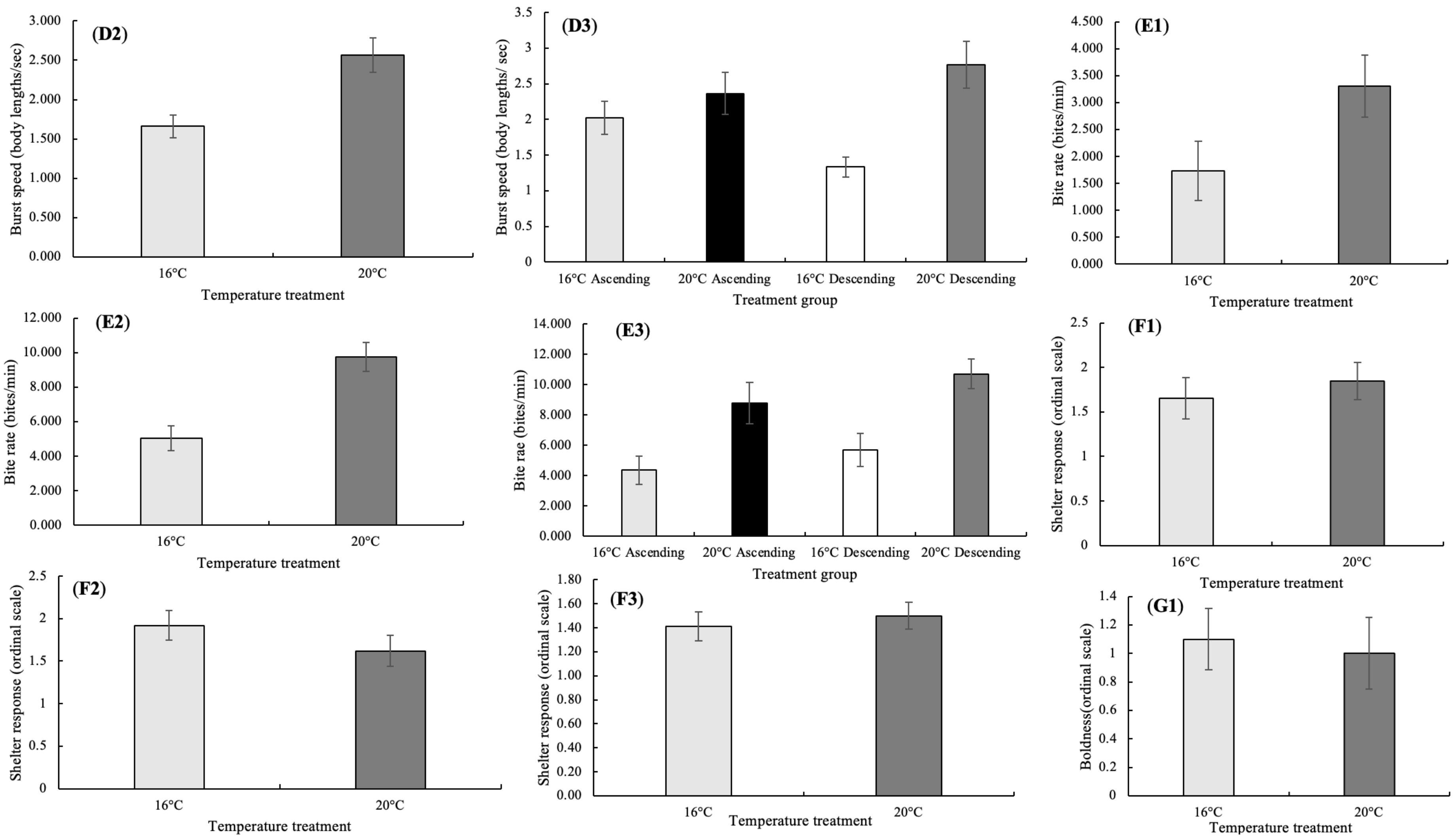
(D2)
Burst speed (body lengths/sec)
3.000
2.500
2.000
1.500
1.000
0.500
0.000
16°C
20°C
Temperature treatment
(D3)
Burst speed (body lengths/ sec)
3.5
3
2.5
2
1.5
1
0.5
0
16°C Ascending
20°C Ascending
16°C Descending
20°C Descending
Treatment group
(E1)
Bite rate (bites/min)
4.500
4.000
3.500
3.000
2.500
2.000
1.500
1.000
0.500
0.000
16°C
20°C
Temperature treatment
(E2)
Bite rate (bites/min)
12.000
10.000
8.000
6.000
4.000
2.000
0.000
16°C
20°C
Temperature treatment
(E3)
Bite rae (bites/min)
14.000
12.000
10.000
8.000
6.000
4.000
2.000
0.000
16°C Ascending
20°C Ascending
16°C Descending
20°C Descending
Treatment group
(F1)
Shelter response (ordinal scale)
2.5
2
1.5
1
0.5
0
16°C
20°C
Temperature treatment
(F2)
Shelter response (ordinal scale)
2.5
2
1.5
1
0.5
0
16°C
20°C
Temperature treatment
(F3)
Shelter response (ordinal scale)
1.80
1.60
1.40
1.20
1.00
0.80
0.60
0.40
0.20
0.00
16°C
20°C
Temperature treatment
(G1)
Boldness(ordinal scale)
1.4
1.2
1
0.8
0.6
0.4
0.2
0
16°C
20°C
Temperature treatment

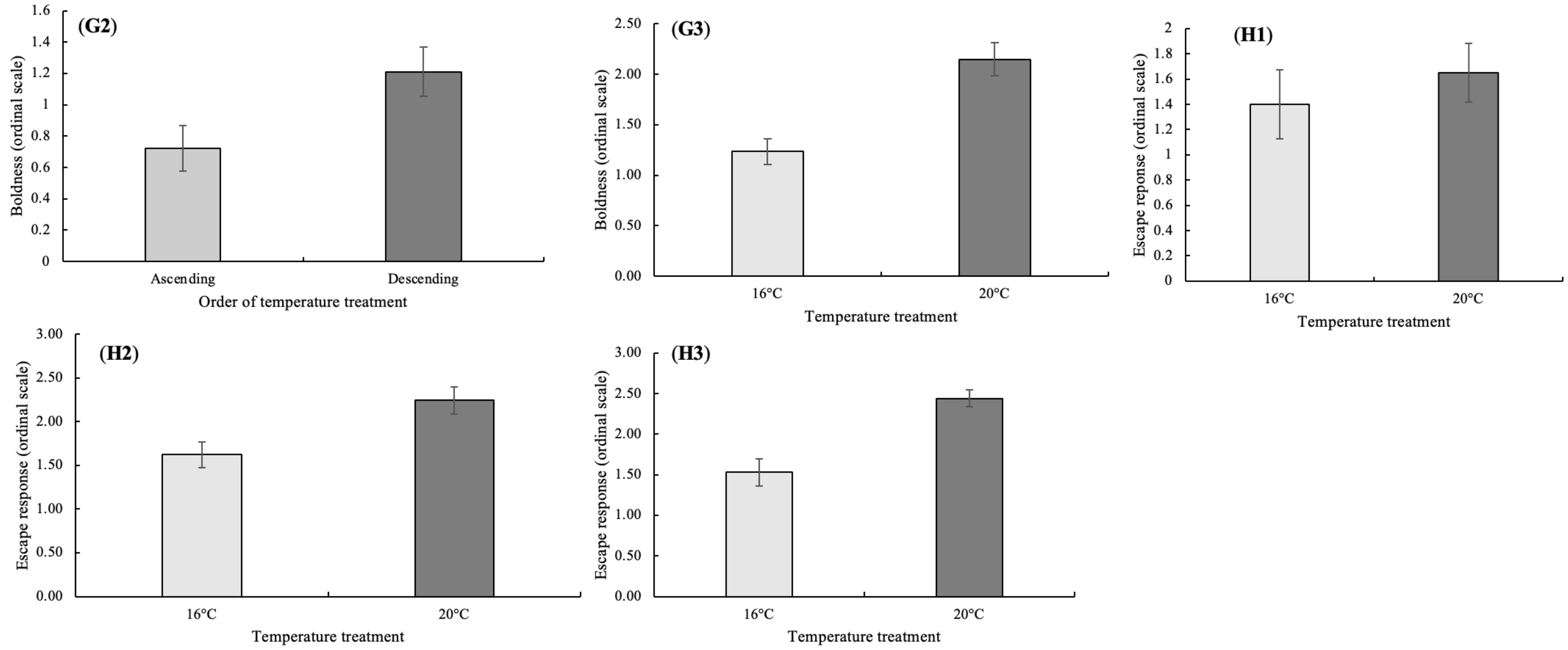


**Figure S2.** Various performance metrics averages across treatments (mean ± SEM) across species (*Centropogon australis, Gerres subfasciatus* and *Pelates sexlineatus*). **(A1)** Average instantaneous growth rate ($G_{inst}$, g/day) in *C. australis* ($n_{16}$= 18, $n_{20}$=19); **(A2)** Average instantaneous growth rate ($G_{inst}$, g/day) in *G. subfasciatus* ($n_{16}$=35, $n_{20}$=36); **(A3)** Average instantaneous growth rate ($G_{inst}$, g/day) in *P. sexlineatus* ($n_{16Asc}$=17, $n_{20Asc}$=18, $n_{16Des}$=15, $n_{20Des}$=15); **(B1)** Average change in total length (mm/day) in *C. australis* ($n_{16}$= 20, $n_{20}$=19); **(B2)** Average change in total length (mm/day) in *G. subfasciatus* ($n_{16}$= 33, $n_{20}$=33); **(B3)** Average change in total length (mm/day) in *P. sexlineatus* ($n_{16Asc}$=18, $n_{20Asc}$=15, $n_{16Des}$=15, $n_{20Des}$=18); **(C1)** Average aerobic scope (mg $O_2$ $kg^{-1}$ $h^{-1}$) in *G. subfasciatus* ($n_{16}$= 37, $n_{20}$=37); **(C2)** Average aerobic scope (mg $O_2$ $kg^{-1}$ $h^{-1}$) in *P. sexlineatus* ($n_{16}$=34, $n_{20}$=33); **(D1)** Average burst speed (body lengths/sec) in *C. australis* ($n_{16}$= 19, $n_{20}$=17); **(D2)** Average burst speed (body lengths/sec) in *G. subfasciatus* ($n_{16}$= 36, $n_{20}$=36); **(D3)** Average burst speed (body lengths/sec) in *P. sexlineatus*

*sexlineatus* (n16Asc=17, n20Asc=18, n16Des=19, n20Des=18); **(E1)** Average bite rate (bites/min) in *C. australis* (n16= 20, n20=19); **(E2)** Average bite rate (bites/min) in *G. subfasciatus* (n16= 37, n20=37); **(E3)** Average bite rate (bites/min) in *P. sexlineatus* (n16Asc=18, n20Asc=18, n16Des=19, n20Des=19); **(F1)** Average shelter response (ordinal scale) in *C. australis* (n16= 20, n20=20); **(F2)** Average shelter response (ordinal scale) in *G. subfasciatus* (n16= 37, n20=37); **(F3)** Average shelter response (ordinal scale) in *P. sexlineatus* (n16= 34, n20=34); **(G1)** Average boldness (ordinal scale) in *C. australis* (n16= 20, n20=20); **(G2)** Average boldness (ordinal scale) in *G. subfasciatus* (nAsc= 36, nDes=38); **(G3)** Average boldness (ordinal scale) in *P. sexlineatus* (n16= 34, n20=33); **(H1)** Average escape response (ordinal scale) in *C. australis* (n16= 20, n20=20); **(H2)** Average escape response (ordinal scale) in *G. subfasciatus* (n16= 37, n20=37); **(H3)** Average escape response (ordinal scale) in *P. sexlineatus* (n16= 34, n20=34).

**Table S1 (next page).** ANOVA output tables for differences in performance across species (*Centropogon australis, Gerres subfasciatus* and *Pelates sexlineatus*) with temperature, order of temperatures and food regimes (*Gerres subfasciatus* and *Pelates sexlineatus* only) as factors **(A1)** *C. australis* instantaneous growth rate; **(A2)** *G. subfasciatus* instantaneous growth rate; **(A3)** *P. sexlineatus* instantaneous growth rate; **(B1)** *C. australis* change in total length; **(B2)** *G. subfasciatus* change in total length; **(B3)** *P. sexlineatus* change in total length; **(C1)** *G. subfasciatus* aerobic scope; **(C2)** *P. sexlineatus* aerobic scope; **(D1)** *C. australis* burst speed; **(D2)** *G. subfasciatus* burst speed; **(D3)** *P. sexlineatus* burst speed; **(E1)** *C. australis* bite rate; **(E2)** *G. subfasciatus* bite rate; **(E3)** *P. sexlineatus* bite rate; (**F1**) *C. australis* boldness; **(F2)** *G. subfasciatus* boldness; (**F3**) *P. sexlineatus* boldness; **(G1)** *C. australis* shelter response; **(G2)** *G. subfasciatus* shelter response; **(G3)** *P. sexlineatus* shelter response; **(H1)** *C.australis* escape response; **(H2)** *G. subfasciatus* escape response; **(H3)** *P. sexlineatus* escape response.

**(A1)**

| Effect | Degrees of freedom (df) | Mean square | F | p-value |
|---|---|---|---|---|
| **Temperature** | 1 | 0.002 | 6.84 | 0.013 |
| **Order of temperatures** | 1 | 0.001 | 2.31 | 0.138 |
| **Temperature * Order of temperatures** | 1 | $5.05 \times 10^{-6}$ | 0.019 | 0.892 |
| **Error** | 33 | | | |
| **Total** | 36 | | | |

**(A2)**

| Effect | Degrees of freedom (df) | Mean square | F | p-value |
|---|---|---|---|---|
| **Temperature** | 1 | 0.003 | 27.97 | <0.001 |
| **Order of temperatures** | 1 | 0.000 | 2.67 | 0.107 |
| **Food regime** | 1 | $2.42 \times 10^{-5}$ | 0.26 | 0.612 |
| **Temperature * Order of temperatures** | 1 | $1.97 \times 10^{-5}$ | 0.211 | 0.647 |
| **Temperature * Food regime** | 1 | $5.49 \times 10^{-5}$ | 0.590 | 0.445 |
| **Order of temperatures * Food regime** | 1 | 0.000 | 2.083 | 0.154 |
| **Temperature * Order of temperatures * Food regimes** | 1 | $1.02 \times 10^{-6}$ | 0.011 | 0.917 |
| **Error** | 63 | | | |
| **Total** | 70 | | | |

**(A3)**

| Effect | Degrees of freedom (df) | Mean square | F | p-value |
|---|---|---|---|---|
| **Temperature** | 1 | 0.000 | 10.76 | 0.002 |
| **Order of temperatures** | 1 | 0.000 | 12.53 | <0.001 |
| **Food regime** | 1 | $1.29 \times 10^{-5}$ | 0.704 | 0.405 |
| **Temperature * Order of temperatures** | 1 | 0.001 | 39.82 | <0.001 |
| **Temperature * Food regime** | 1 | $1.53 \times 10^{-6}$ | 0.084 | 0.773 |
| **Order of temperatures * Food regime** | 1 | $1.30 \times 10^{-5}$ | 0.713 | 0.402 |
| **Temperature * Order of temperatures * Food regimes** | 1 | $3.14 \times 10^{-5}$ | 1.72 | 0.195 |
| **Error** | 57 | | | |
| **Total** | 65 | | | |

**(B1)**

| Effect | Degrees of freedom (df) | Mean square | F | p-value |
|---|---|---|---|---|
| **Temperature** | 1 | 8.85 | 4.62 | 0.039 |
| **Order of temperatures** | 1 | 0.063 | 1.026 | 0.318 |
| **Temperature * Order of temperatures** | 1 | $5.05 \times 10^{-6}$ | 0.019 | 0.892 |
| **Error** | 35 | | | |
| **Total** | 39 | | | |

**(B2)**

| Effect | Degrees of freedom (df) | Mean square | F | p-value |
|---|---|---|---|---|
| **Temperature** | 1 | 0.209 | 20.00 | < 0.001 |
| **Order of temperatures** | 1 | 0.007 | 0.705 | 0.404 |
| **Food regime** | 1 | 0.001 | 0.132 | 0.717 |
| **Temperature * Order of temperatures** | 1 | 0.006 | 0.587 | 0.447 |
| **Temperature * Food regime** | 1 | 0.11 | 1.04 | 0.312 |
| **Order of temperatures * Food regime** | 1 | 0.001 | 0.053 | 0.819 |
| **Temperature * Order of temperatures * Food regimes** | 1 | 0.006 | 0.557 | 0.458 |
| **Error** | 58 | | | |
| **Total** | 66 | | | |

**(B3)**

| Effect | Degrees of freedom (df) | Mean square | F | p-value |
|---|---|---|---|---|
| **Temperature** | 1 | 0.234 | 14.86 | <0.001 |
| **Order of temperatures** | 1 | 0.763 | 12.38 | 0.001 |
| **Food regime** | 1 | 0.004 | 0.276 | 0.601 |
| **Temperature * Order of temperatures** | 1 | 0.377 | 23.87 | <0.001 |
| **Temperature * Food regime** | 1 | 0.040 | 2.57 | 0.114 |
| **Order of temperatures * Food regime** | 1 | 0.002 | 0.113 | 0.738 |
| **Temperature * Order of temperatures * Food regimes** | 1 | 0.024 | 1.54 | 0.219 |
| **Error** | 58 | | | |
| **Total** | 66 | | | |

**(C1)**

| Effect | Degrees of freedom (df) | Mean square | F | p-value |
|---|---|---|---|---|
| **Temperature** | 1 | 3.72 | 39.72 | <0.001 |
| **Order of temperatures** | 1 | 0.0006 | 0.065 | 0.800 |
| **Food regime** | 1 | 0.154 | 1.640 | 0.205 |
| **Temperature * Order of temperatures** | 1 | 0.239 | 0.277 | 0.600 |
| **Temperature * Food regime** | 1 | 0.496 | 0.574 | 0.451 |
| **Order of temperatures * Food regime** | 1 | 0.089 | 0.103 | 0.749 |
| **Temperature * Order of temperatures * Food regimes** | 1 | 0.005 | 0.064 | 0.801 |
| **Error** | 58 | | | |
| **Total** | 66 | | | |

**(C2)**

| Effect | Degrees of freedom (df) | Mean square | F | p-value |
|---|---|---|---|---|
| **Temperature** | 1 | 94848.68 | 6.99 | 0.01 |
| **Order of temperatures** | 1 | 5544.251 | 0.409 | 0.525 |
| **Food regime** | 1 | 59980.13 | 3.535 | 0.065 |
| **Temperature * Order of temperatures** | 1 | 481.063 | 0.035 | 0.851 |
| **Temperature * Food regime** | 1 | 20378.64 | 1.503 | 0.225 |
| **Order of temperatures * Food regime** | 1 | 19032.027 | 1.403 | 0.241 |
| **Temperature * Order of temperatures * Food regimes** | 1 | 15097.518 | 1.113 | 0.296 |
| **Error** | 59 | | | |
| **Total** | 67 | | | |

**(D1)**

| Effect | Degrees of freedom (df) | Mean square | F | p-value |
|---|---|---|---|---|
| **Temperature** | 1 | 0.773 | 1.55 | 0.223 |
| **Order of temperatures** | 1 | 0.024 | 0.048 | 0.828 |
| **Temperature * Order of temperatures** | 1 | 1.54 | 3.09 | 0.089 |
| **Error** | 35 | | | |
| **Total** | 39 | | | |

**(D2)**

| Effect | Degrees of freedom (df) | Mean square | F | p-value |
|---|---|---|---|---|
| **Temperature** | 1 | 13.88 | 11.24 | 0.001 |
| **Order of temperatures** | 1 | 0.355 | 0.288 | 0.594 |
| **Food regime** | 1 | 0.032 | 0.026 | 0.873 |
| **Temperature * Order of temperatures** | 1 | 1.44 | 0.062 | 0.804 |
| **Temperature * Food regime** | 1 | 0.230 | 0.187 | 0.667 |
| **Order of temperatures * Food regime** | 1 | 0.818 | 0.663 | 0.419 |
| **Temperature * Order of temperatures * Food regimes** | 1 | 0.062 | 0.050 | 0.823 |
| **Error** | 64 | | | |
| **Total** | 72 | | | |

**(D3)**

| Effect | Degrees of freedom (df) | Mean square | F | p-value |
|---|---|---|---|---|
| Temperature | 1 | 18.95 | 12.45 | <0.001 |
| Order of temperatures | 1 | 25.04 | 16.45 | <0.001 |
| Food regime | 1 | 0.865 | 0.586 | 0.454 |
| Temperature * Order of temperatures | 1 | 47.06 | 30.92 | <0.001 |
| Temperature * Food regime | 1 | 0.367 | 0.241 | 0.626 |
| Order of temperatures * Food regime | 1 | 1.14 | 0.747 | 0.391 |
| Temperature * Order of temperatures * Food regimes | 1 | 0.246 | 0.162 | 0.689 |
| Error | 64 | | | |
| Total | 72 | | | |

**(E1)**

| Effect | Degrees of freedom (df) | Mean square | F | p-value |
|---|---|---|---|---|
| Temperature | 1 | 0.773 | 1.55 | 0.223 |
| Order of temperatures | 1 | 0.024 | 0.048 | 0.828 |
| Temperature * Order of temperatures | 1 | 1.542 | 3.086 | 0.089 |
| Error | 32 | | | |
| Total | 36 | | | |

**(E2)**

| Effect | Degrees of freedom (df) | Mean square | F | p-value |
|---|---|---|---|---|
| Temperature | 1 | 408.69 | 17.58 | <0.001 |
| Order of temperatures | 1 | 49.45 | 2.13 | 0.149 |
| Food regime | 1 | 14.67 | 0.631 | 0.430 |
| Temperature * Order of temperatures | 1 | 1.444 | 0.062 | 0.804 |
| Temperature * Food regime | 1 | 0.194 | 0.008 | 0.927 |
| Order of temperatures * Food regime | 1 | 17.879 | 0.769 | 0.384 |
| Temperature * Order of temperatures * Food regimes | 1 | 0.624 | 0.027 | 0.870 |
| Error | 66 | | | |
| Total | 74 | | | |

**(E3)**

| Effect | Degrees of freedom (df) | Mean square | F | p-value |
|---|---|---|---|---|
| Temperature | 1 | 734.28 | 8.41 | 0.005 |
| Order of temperatures | 1 | 1597.26 | 18.29 | <0.001 |
| Food regime | 1 | 93.74 | 1.073 | 0.304 |
| Temperature * Order of temperatures | 1 | 973.78 | 11.15 | 0.001 |
| Temperature * Food regime | 1 | 105.63 | 1.21 | 0.276 |
| Order of temperatures * Food regime | 1 | 6.48 | 0.074 | 0.786 |
| Temperature * Order of temperatures * Food regimes | 1 | 33.46 | 0.383 | 0.538 |
| Error | 60 | | | |
| Total | 68 | | | |

**(F1)**

| Effect | Degrees of freedom (df) | Mean square | F | p-value |
|---|---|---|---|---|
| Temperature | 1 | 0.100 | 0.098 | 0.756 |
| Order of temperatures | 1 | 1.600 | 1.574 | 0.218 |
| Temperature * Order of temperatures | 1 | 3.600 | 3.541 | 0.068 |
| Error | 36 | | | |
| Total | 40 | | | |

**(F2)**

| Effect | Degrees of freedom (df) | Mean square | F | p-value |
|---|---|---|---|---|
| Temperature | 1 | 0.447 | 0.451 | 0.504 |
| Order of temperatures | 1 | 4.719 | 4.76 | 0.033 |
| Food regime | 1 | 1.313 | 1.324 | 0.254 |
| Temperature * Order of temperatures | 1 | 0.082 | 0.083 | 0.775 |
| Temperature * Food regime | 1 | 1.313 | 1.324 | 0.254 |
| Order of temperatures * Food regime | 1 | 2.736 | 2.736 | 0.103 |
| Temperature * Order of temperatures * Food regimes | 1 | 0.036 | 0.037 | 0.849 |
| Error | 66 | | | |
| Total | 74 | | | |

**(F3)**

| Effect | Degrees of freedom (df) | Mean square | F | p-value |
|---|---|---|---|---|
| Temperature | 1 | 10.81 | 16.14 | <0.001 |
| Order of temperatures | 1 | 0.040 | 0.060 | 0.808 |
| Food regime | 1 | 0.511 | 0.763 | 0.386 |
| Temperature * Order of temperatures | 1 | 0.099 | 0.148 | 0.702 |
| Temperature * Food regime | 1 | 0.360 | 0.538 | 0.466 |
| Order of temperatures * Food regime | 1 | 0.003 | 1.748 | 0.191 |
| Temperature * Order of temperatures * Food regimes | 1 | 0.007 | 0.011 | 0.917 |
| Error | 60 | | | |
| Total | 68 | | | |

**(G1)**

| Effect | Degrees of freedom (df) | Mean square | F | p-value |
|---|---|---|---|---|
| Temperature | 1 | 0.100 | 0.098 | 0.756 |
| Order of temperatures | 1 | 1.600 | 1.574 | 0.218 |
| Temperature * Order of temperatures | 1 | 3.600 | 3.541 | 0.068 |
| Error | 36 | | | |
| Total | 40 | | | |

**(G2)**

| Effect | Degrees of freedom (df) | Mean square | F | p-value |
|---|---|---|---|---|
| Temperature | 1 | 1.950 | 1.922 | 0.170 |
| Order of temperatures | 1 | 1.755 | 1.730 | 0.193 |
| Food regime | 1 | 1.368 | 1.380 | 0.244 |
| Temperature * Order of temperatures | 1 | 1.313 | 1.324 | 0.254 |
| Temperature * Food regime | 1 | 0.195 | 0.192 | 0.662 |
| Order of temperatures * Food regime | 1 | 1.128 | 1.112 | 0.295 |
| Temperature * Order of temperatures * Food regimes | 1 | 2.155 | 2.125 | 0.150 |
| Error | 66 | | | |
| Total | 74 | | | |

**(G3)**

| Effect | Degrees of freedom (df) | Mean square | F | p-value |
|---|---|---|---|---|
| Temperature | 1 | 0.511 | 0.763 | 0.386 |
| Order of temperatures | 1 | 0.206 | 0.341 | 0.561 |
| Food regime | 1 | 0.931 | 1.540 | 0.220 |
| Temperature * Order of temperatures | 1 | 0.003 | 0.005 | 0.942 |
| Temperature * Food regime | 1 | 0.116 | 0.192 | 0.663 |
| Order of temperatures * Food regime | 1 | 0.128 | 0.200 | 0.656 |
| Temperature * Order of temperatures * Food regimes | 1 | 0.029 | 0.048 | 0.827 |
| Error | 59 | | | |
| Total | 67 | | | |

(H1)

| Effect | Degrees of freedom (df) | Mean square | F | p-value |
|---|---|---|---|---|
| Temperature | 1 | 0.626 | 0.474 | 0.496 |
| Order of temperatures | 1 | 1.225 | 0.928 | 0.342 |
| Temperature * Order of temperatures | 1 | 0.625 | 0.474 | 0.496 |
| Error | 36 | | | |
| Total | 40 | | | |

(H2)

| Effect | Degrees of freedom (df) | Mean square | F | p-value |
|---|---|---|---|---|
| Temperature | 1 | 6.957 | 8.051 | 0.006 |
| Order of temperatures | 1 | 1.571 | 1.817 | 0.182 |
| Food regime | 1 | 0.004 | 0.004 | 0.949 |
| Temperature * Order of temperatures | 1 | 0.239 | 0.277 | 0.600 |
| Temperature * Food regime | 1 | 0.496 | 0.574 | 0.451 |
| Order of temperatures * Food regime | 1 | 0.089 | 0.103 | 0.749 |
| Temperature * Order of temperatures * Food regimes | 1 | 2.086 | 2.414 | 0.125 |
| Error | 66 | | | |
| Total | 74 | | | |

(H3)

| Effect | Degrees of freedom (df) | Mean square | F | p-value |
|---|---|---|---|---|
| Temperature | 1 | 19.00 | 34.07 | <0.001 |
| Order of temperatures | 1 | 0.311 | 0.557 | 0.458 |
| Food regime | 1 | 0.280 | 0.501 | 0.482 |
| Temperature * Order of temperatures | 1 | 0.099 | 0.148 | 0.702 |
| Temperature * Food regime | 1 | 0.128 | 0.229 | 0.634 |
| Order of temperatures * Food regime | 1 | 1.691 | 3.033 | 0.087 |
| Temperature * Order of temperatures * Food regimes | 1 | 0.01 | 0.018 | 0.894 |
| Error | 60 | | | |
| Total | 68 | | | |

**Table S2 (next page).** Outputs regressions between performance metrics across species. **(A)** *Centropogon australis* bite rate (bites/min) vs change in total length (mm/day), $R^2$ = 0.12; **(B)** *Gerres subfasciatus* instantaneous growth rate ($G_{inst}$, g/day) vs change in total length (mm/day), $R^2$ = 0.18; **(C)** *G. subfasciatus* bite rate (bites/min) vs instantaneous growth rate ($G_{inst}$, g/day), $R^2$ = 0.08; **(D)** *G. subfasciatus* aerobic scope (mg $O_2$/ kg h) vs instantaneous growth rate ($G_{inst}$, g/day), $R^2$ = 0.2; **(E)** *G. subfasciatus* aerobic scope (mg $O_2$/ kg h) vs change in total length (mm/day), $R^2$ = 0.07; **(F)** *G. subfasciatus* bite rate (bites/min) vs aerobic scope (mg $O_2$/ kg h), $R^2$ = 0.08; **(G)** *G. subfasciatus* aerobic scope (mg $O_2$/ kg h) vs burst speed (body lengths/sec), $R^2$ = 0.07; **(H)** *G. subfasciatus* burst speed (body lengths/sec) vs escape response (ordinal scale), McFadden $R^2$ = 0.13; **(I)** *G. subfasciatus* boldness (ordinal scale) vs escape response (ordinal scale), McFadden $R^2$ = 0.09; **(J)** *Pelates sexlineatus* instantaneous growth rate ($G_{inst}$, g/day) vs bite rate (bites/min), $R^2$ = 0.27; **(K)** *P. sexlineatus* instantaneous growth rate ($G_{inst}$, g/day) vs change in total length (mm/day), $R^2$ = 0.20; **(L)** *P. sexlineatus* aerobic scope (mg $O_2$/ kg h) vs burst speed (body lengths/sec), $R^2$ = 0.06; **(M)** *P. sexlineatus* aerobic scope (mg $O_2$/ kg h) vs boldness (ordinal scale), McFadden $R^2$ = 0.08; **(N)** *P. sexlineatus* aerobic scope (mg $O_2$/ kg h) vs escape response (ordinal scale), McFadden $R^2$ = 0.08; **(O)** *P. sexlineatus* burst speed (body lengths/sec) vs escape response (ordinal scale), McFadden $R^2$ = 0.24; **(P)** *P. sexlineatus* boldness (ordinal scale) vs bite rate (bites/min), McFadden $R^2 = 0.15$; **(Q)** *P. sexlineatus* boldness (ordinal scale) vs burst speed (body lengths/sec), McFadden $R^2 =$ 0.08; **(R)** *P. sexlineatus* boldness (ordinal scale) vs escape response (ordinal scale), McFadden $R^2 = 0.12$.

(A)

| Model | Degrees of freedom (df) | F | p-value |
|---|---|---|---|
| Regression | 1 | 4.58 | 0.039 |
| Residual | 36 | | |
| Total | 37 | | |

(B)

| Model | Degrees of freedom (df) | F | p-value |
|---|---|---|---|
| Regression | 1 | 14.02 | < 0.001 |
| Residual | 63 | | |
| Total | 64 | | |

(C)

| Model | Degrees of freedom (df) | F | p-value |
|---|---|---|---|
| Regression | 1 | 5.84 | 0.0018 |
| Residual | 69 | | |
| Total | 70 | | |

(D)

| Model | Degrees of freedom (df) | F | p-value |
|---|---|---|---|
| Regression | 1 | 17.36 | <0.001 |
| Residual | 69 | | |
| Total | 70 | | |

(E)

| Model | Degrees of freedom (df) | F | p-value |
|---|---|---|---|
| Regression | 1 | 5.25 | 0.025 |
| Residual | 66 | | |
| Total | 67 | | |

(F)

| Model | Degrees of freedom (df) | F | p-value |
|---|---|---|---|
| Regression | 1 | 5.92 | 0.017 |
| Residual | 72 | | |
| Total | 73 | | |

(G)

| Model | Degrees of freedom (df) | F | p-value |
|---|---|---|---|
| Regression | 1 | 5.45 | 0.022 |
| Residual | 70 | | |
| Total | 71 | | |

(H)

| Chi-Square | Degrees of freedom (df) | p-value |
|---|---|---|
| 183.14 | 71 | < 0.001 |

(I)

| Chi-Square | Degrees of freedom (df) | p-value |
|---|---|---|
| 16.29 | 3 | < 0.001 |

(J)

| Model | Degrees of freedom (df) | F | p-value |
|---|---|---|---|
| Regression | 1 | 23.82 | < 0.001 |
| Residual | 64 | | |
| Total | 65 | | |

(K)

| Model | Degrees of freedom (df) | F | p-value |
|---|---|---|---|
| Regression | 1 | 15.98 | < 0.001 |
| Residual | 63 | | |
| Total | 64 | | |

(L)

| Model | Degrees of freedom (df) | F | p-value |
|---|---|---|---|
| Regression | 1 | 4.08 | 0.048 |
| Residual | 62 | | |
| Total | 63 | | |

(M)

| Chi-Square | Degrees of freedom (df) | p-value |
|---|---|---|
| 164.18 | 66 | < 0.001 |

(N)

| Chi-Square | Degrees of freedom (df) | p-value |
|---|---|---|
| 171.57 | 66 | < 0.001 |

(O)

| Chi-Square | Degrees of freedom (df) | p-value |
|---|---|---|
| 166.79 | 64 | < 0.001 |

(P)

| Chi-Square | Degrees of freedom (df) | p-value |
|---|---|---|
| 157.08 | 61 | < 0.001 |

(Q)

| Chi-Square | Degrees of freedom (df) | p-value |
|---|---|---|
| 158.98 | 64 | < 0.001 |

(R)

| Chi-Square | Degrees of freedom (df) | p-value |
|---|---|---|
| 20.12 | 3 | < 0.001 |

**Fish laboratory growth vs. growth in the wild – an otolith analysis**

Otolith increment growth width (proxy of fish growth) in the wild did not differed to the one in laboratory conditions for both *Centropogon australis* ($t(9)= -0.60$ , $p = 0.57$ ) and *Pelates sexlineatus* ($t(8) = 0.09$ , $p = 0.92$). The increment growth rate in the wild was a good predictor of the increment growth rate in the lab.

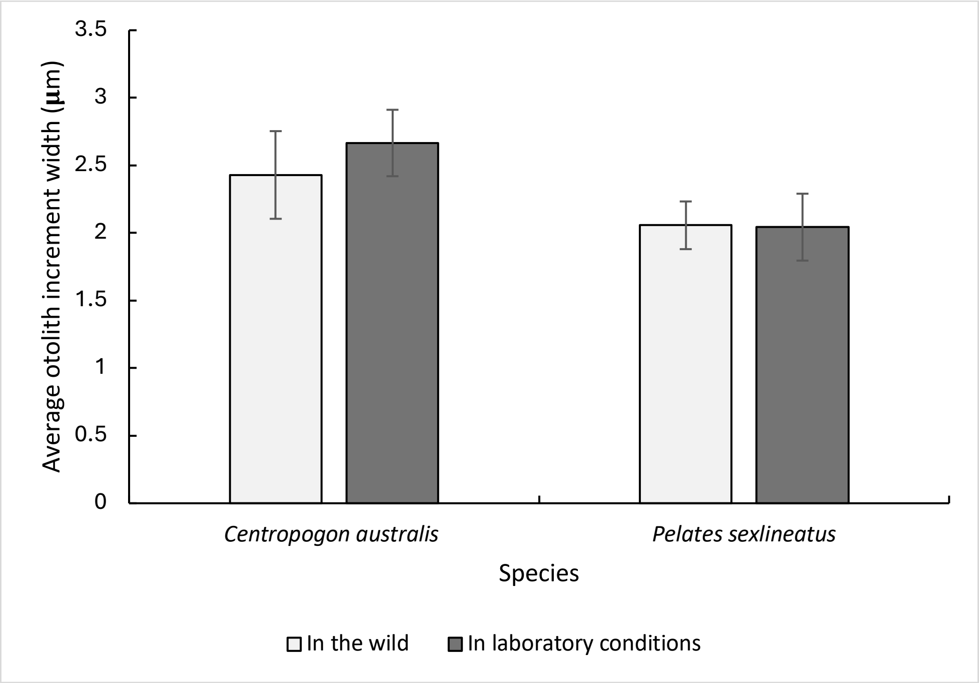


**Figure S3.** Average otoliths increment width (µm) in the wild vs in laboratory conditions in *Centropogon australis* and *Pelates sexlineatus* (mean ± SEM). *Centropogon australis* average otolith increment width (µm) in the wild at approximately 16°C (2.43 ± 0.32) vs average otolith increment width in laboratory conditions at 16°C (2.67 ± 0.25). (B) *Pelates sexlineatus* average otolith increment width (µm) in the wild at approximately 20°C (2.06 ± 0.18) vs average otolith increment width in laboratory conditions at 20°C (2.04 ± 0.25). Each point represents one fish otolith growth in lab vs field.
